# parafac4microbiome: Exploratory analysis of longitudinal microbiome data using Parallel Factor Analysis

**DOI:** 10.1101/2024.05.02.592191

**Authors:** G.R. van der Ploeg, J.A. Westerhuis, A. Heintz-Buschart, A.K. Smilde

## Abstract

Studies investigating microbial temporal dynamics are increasingly common, leveraging longitudinal designs that collect microbial abundance data across multiple time points from the same subjects. Traditional exploratory approaches like Principal Component Analysis (PCA) fail to fully utilize this structure. By organizing data as a three-way array—subjects as rows, microbial abundances as columns, and time points as the third dimension—multi-way methods such as Parallel Factor Analysis (PARAFAC) can better capture temporal and structural patterns. This study demonstrates Parallel Factor Analysis (PARAFAC) as a method to explore longitudinal microbiome data using three exemplary studies.

In the first example, a long time series of in vitro microbiomes, PARAFAC identifies primary time-resolved variations. The second example, a longitudinal infant gut microbiome study, shows that PARAFAC can distinguish subject groups and enhance comparative analysis, even with moderate missing data. In the third example, a gingivitis intervention study of the oral microbiome, PARAFAC enables the identification of microbial subcommunities of interest through post-hoc clustering.

These examples highlight PARAFAC’s broad applicability for analysing longitudinal microbiome data across diverse environments. The approach is implemented in the R package *parafac4microbiome*, available on CRAN, providing researchers with accessible tools for similar analyses.

**Importance:** Understanding how microbiomes change over time can give us valuable insights into their role in health and disease. Many traditional methods like Principal Component Analysis (PCA) miss important patterns in data collected over time, but Parallel Factor Analysis (PARAFAC) helps uncover these trends in a much clearer way. Using this approach, we were able to identify key changes in microbiomes across different settings, like lab experiments, the infant gut, and the mouth. PARAFAC also works well even when some data is missing, which is a common issue. To make this tool accessible, we have included it in a user-friendly R package, enabling other researchers to analyse microbiome dynamics in their own studies and explore how these changes might influence health and treatments.

## Background

The role of the microbiome in biological systems and in disease has been thoroughly explored [1– 3]. However, most studies are cross-sectional while microbial temporal dynamics are required to understand host-microbiome and microbe-microbe interactions [4, 5]. Recently, studies that investigate these dynamics have become more frequent [6–8].

In a longitudinal microbiome study design, microbial abundance data are collected across multiple time points from the same subjects [9–11]. These data are often stored in a two-way data table, where the rows constitute the measurements (samples from multiple individuals over time), and the columns contain the microbial abundances (**Figure 1A**). Two-way ordination methods like Principal Component Analysis (PCA) [12, 13] or Multidimensional Scaling (MDS) [14] are then employed to explore the data (**Figure 1B**). However, this often results in hard-to-interpret visualizations because these methods treat every measurement as a separate object to be ordinated (**Figure 1D, E**). As such, there is a need for exploratory methods that take the crossed longitudinal study design into account.

**Figure 1.**
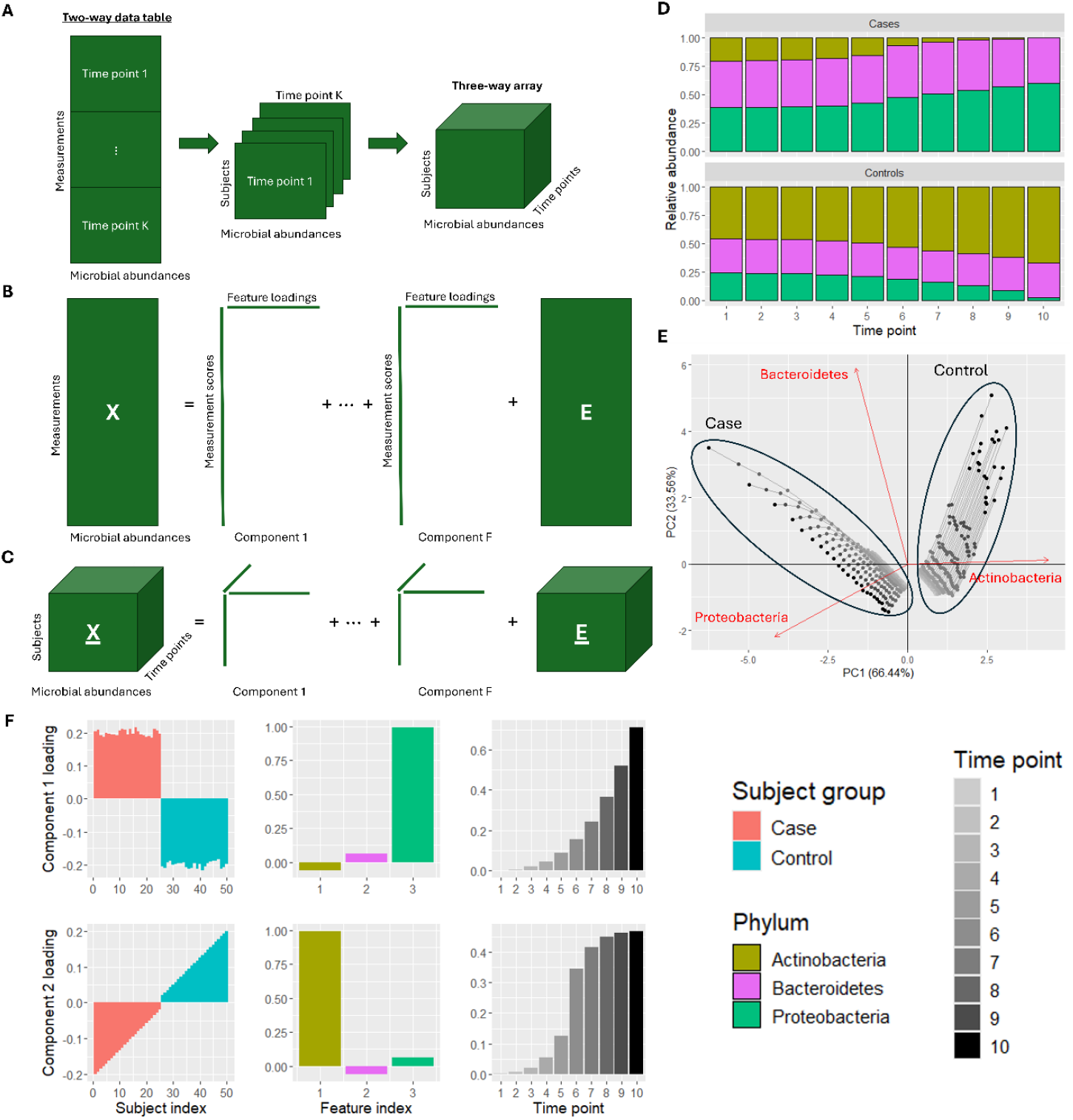
(A) Longitudinal microbiome data is often presented as a two-way data table with measurements (i.e. one sample for every subject-time point combination) in the rows and microbial abundances in the columns. This kind of data can be converted to a three-way array by putting all measurements for each time point in the third dimension. (B) Principal Component Analysis decomposes a two-way data table into components with scores for the measurements and loadings for the microbial relative abundances. This does not take the longitudinal study design into account. (C) Parallel factor analysis decomposes a three-way data array into components with three loading vectors each: one for the subjects, one for the microbial relative abundances, and one for time. (D) In a mock intervention dataset, we have obtained count data of three phyla (Actinobacteria, Bacteroidetes and Proteobacteria) corresponding to 25 case and 25 control subjects across 10 time points. At time point 1, all subjects look very similar to each other. At time point 10 the case subjects are depleted of Actinobacteria and enriched in Proteobacteria, while for the control subjects the opposite is true. The magnitude of the depletion and enrichment of Actinobacteria at time point 10 varies substantially within the case and control groups, respectively. The simulated counts corresponding to Bacteroidetes have no time-resolved subject group differences. (E) Biplot of a Principal Component Analysis of the mock dataset after centred log-ratio transformation, centring and scaling. Samples taken from the same subject are connected with a line. The first principal component, explaining 66.44% of the variation, separates the case and control samples based on the abundance of Actinobacteria. The second principal component, explaining 33.56% of the variation, separates the time points using the Proteobacteria and Bacteroidetes abundances. However, the time profiles per subject group are unclear. (F) Plot of the Parallel Factor Analysis model of the data, describing subject group differences and their time profiles. In component 1 the exponential growth of Proteobacteria in the case subjects is identified. In component 2 the sigmoid shaped growth curve of Actinobacteria in the control subjects is identified. The magnitude of the depletion and enrichment of Actinobateria within the case and control groups is visible as a gradient of loading values. Bacteroidetes has a loading close to zero in both components, correctly identifying it as having no discriminating time-resolved variation. This illustrates that PARAFAC can capture various time-resolved differences in microbial abundances between the subject groups in a more interpretable way than PCA.

The longitudinal study design can be used to create a three-way data array where the rows constitute the subjects, the columns contain the microbial abundances, and the third dimension contains the time points (**Figure 1A**). Three-way ordination methods like Parallel Factor Analysis (PARAFAC; [15, 16]) can then be used to explore the data (**Figure 1C**). By making full use of the study design, the model can identify subject groups as well as their time profiles (**Figure 1D, F**).

In this work we show how Parallel Factor Analysis can be used to explore longitudinal microbiome data. We first give a mathematical description of the PARAFAC method and its main requirement, followed by a brief review of longitudinal microbiome data processing needed to meet this requirement and the principles of PARAFAC model selection. We then highlight the benefits of making a PARAFAC model using three exemplary datasets. In the first example study [6] we highlight the time-resolved variation that PARAFAC models. In the second study [7], we highlight how PARAFAC can find differences between subject groups and enhance comparative analysis despite a moderate amount of missing data. In the third study [8] we highlight a post-hoc clustering approach to identify microbes of interest with PARAFAC. The analyses and the example datasets with the resulting figures are implemented in the R package parafac4microbiome, which is available on CRAN at https://cran.rstudio.com/web/packages/parafac4microbiome/.

## Methods

### Notation and definitions

We briefly define the mathematical notation that will be used throughout this paper. Scalars are indicated by lower-case italics such as *a*. Vectors are indicated by bold lower-case characters such as ***b***. Two-way matrices are indicated with bold capitalized characters such as ***X***. Underlined bold capitalized characters are used for three-way arrays such as *<u>**X**</u>*. The letters *I, J*, and *K* are reserved to indicate the dimension of the subject, microbial abundance, and time mode, respectively. Hence the element for subject *i* ∈ (1 … *I*), microbial abundance *j* ∈ (1 … *J*) at timepoint *k* ∈ (1 … *K*) of a three-way array *<u>**X**</u>* is called *x*_*ijk*_. We create the data cubes of microbial relative abundances such that the subjects are in the first mode, the microbial taxa are in the second mode, and time points are in the third mode. We will not distinguish between the terms ‘factor’ and ‘component’, nor between the terms ‘way’ and ‘mode’. Generally, we will use the words ‘component’ and ‘mode’ throughout this paper.

### Mathematical definition of Parallel Factor Analysis

Parallel Factor Analysis (PARAFAC) [15, 16] is a factor decomposition method that generalizes Principal Component Analysis (PCA) to multi-way data. First, we give a brief mathematical summary of PCA to make the generalizing principle of PARAFAC clearer. We then describe the algorithmic implementation of this method and its properties.

A longitudinal microbiome dataset is often represented as a two-way matrix ***X*** (*IK* × *J*) with all time points for all subjects as rows and the microbial abundances as columns (**Figure 1B**). For convenience, we will refer to a subject-timepoint combination as a measurement *m* ∈ (1 … *M*) where *M* = *IK*. A PCA model of this two-way dataset ***X*** (*M* × *J*) is then given by the two matrices ***T*** (*M* × *F*) and ***P*** (*J* × *F*) containing the scores and loadings of all *F* components respectively. The scores describe the differences between the measurements, and the loadings describe the microbial abundances that are different. A PCA model of such a two-way matrix can be formulated elementwise as

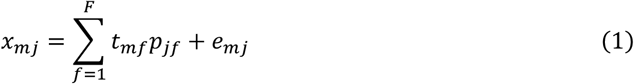

with *x*_*mj*_ the matrix element corresponding to measurement *m* and feature *j, t*_*mf*_ the score of measurement *m* in component *f, p*_*jf*_ the loading of feature *j* in component *f* and *e*_*mj*_ the residual of the matrix element. PCA is a bilinear model as we are using *F* sets of two vectors to approximate the dataset ***X***. This also reveals the main issue at hand: the measurements *M* are ordinated without taking the connection to the subjects *I* and time points *K* into account.

A three-way data array *<u>**X**</u>* is defined as having *I* subjects in the rows, *J* microbial abundances in the columns and *K* time points in the third dimension (**Figure 1C**). A PARAFAC model of a three-way data array *<u>**X**</u>* (*I* × *l* × *K*) is given by three matrices ***A*** (*I* × *F*), ***B*** (*J* × *F*) and ***C*** (*K* × *F*) containing the loadings of all *F* components corresponding to the subject, feature, and time mode, respectively. The element-wise definition of the model is formulated as

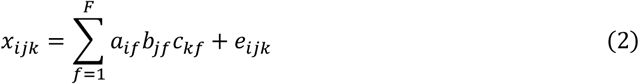

with *x*_*ijk*_ the matrix element corresponding to subject *i*, feature *j* at time point *k, a*_*if*_ the loading of subject *i* of component *f, b*_*jf*_ the loading of feature *j* of component *f* and *c*_*kf*_ the loading of time point *k* of component *f* and *e*_*ijk*_ the residual of the matrix element. In the elementwise definition, the generalization of PCA is readily visible: the loading for the time mode *c*_*kf*_ has been added to the calculation of element *x*_*ijk*_ to represent the time factor. PARAFAC is a trilinear model as we are using *F* sets of three vectors to approximate the data array ***X***.

### Algorithmic implementation of Parallel Factor Analysis

The objective of the algorithmic implementation of PARAFAC is to minimise the sum-of-squares of the residuals through the loss function

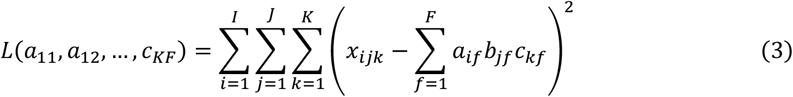

Various implementations of estimating the PARAFAC loading matrices exist [17]. In this paper we estimate the PARAFAC loading matrices through an iterative alternating least-squares algorithm (ALS; [16, 18]), as this implementation is well-documented and robust towards missing values or time points (**Table 1**).

**Table 1:** Algorithmic implementation of the alternating least squares based PARAFAC model estimation.

| Parallel Factor Analysis (PARAFAC) ALS algorithm for a given three-way array $\underline{X}$ | |
| --- | --- |
| (0) | Decide on the number of components $F$ |
| (1) | Initialise $\mathbf{B}$ and $\mathbf{C}$ |
| (2) | Estimate $\mathbf{A}$ from $\underline{X}, \mathbf{B}$ and $\mathbf{C}$ such that $L(a_{11}, a_{12}, \dots, c_{KF})$ is minimised |
| (3) | Estimate $\mathbf{B}$ from $\underline{X}, \mathbf{A}$ and $\mathbf{C}$ such that $L(a_{11}, a_{12}, \dots, c_{KF})$ is minimised |
| (4) | Estimate $\mathbf{C}$ from $\underline{X}, \mathbf{A}$ and $\mathbf{B}$ such that $L(a_{11}, a_{12}, \dots, c_{KF})$ is minimised |
| (5) | Continue to step (2) until convergence. |

Various approaches for initialisation of ***B*** and ***C*** have been proposed and are documented elsewhere [19]. In this paper, we used randomly initialised vectors to determine the appropriate number of components and singular value decomposition-based initialised vectors to investigate the stability of the loadings.

We use the PARAFAC algorithm supplied in the parafac4microbiome R package (v1.1.2, [20]). With our default settings, the model converges in step (5) when the change in sum-of-squares is lower than 1e-4, or terminates after 500 iterations of steps 2-4. In this paper, the convergence threshold is set lower to ensure that PARAFAC models corresponding to a global minimum are reported. Fitting a model for any of the processed datasets in this paper takes less than a minute of computational time on an average computer (Microsoft Windows 11 Home v10.0.22631 with an 12^th^ Gen Intel® Core™ i5-12400F running 2500 MHz with 6 cores or 12 logical processors and 16 Gb RAM).

### PARAFAC model constraints and properties

A PARAFAC decomposition is unique, apart from trivial rescaling and permutations of the factors. This means that estimated component matrices cannot be rotated without a loss of fit. This is a distinct advantage over Principal Component Analysis [16, 21–24]. Additionally, PARAFAC can capture any type of temporal dynamics – linear, exponential or otherwise - because it models time categorically. Thus, the time points are not connected like in a dynamic model. By treating time as a categorical variable, it is assumed that sampling time points are appropriately spaced, and each sampling time point receives a fair contribution to the model. A limitation of PARAFAC is that all subjects must be measured on the same time points. While the method cannot interpolate or extrapolate to time points that were not originally measured, it can interpolate data that is missing in some subjects. Furthermore, the components are orthogonal in PCA while in PARAFAC the components are non-orthogonal by default. The orthogonality constraint in PCA results in models that force interpretation in terms of orthogonal factors which do not always align with the underlying biology, whereas the absence of such a constraint for PARAFAC results in models that can describe processes that are not necessarily independent. Non-orthogonality is an advantage because it is more likely a realistic representation of the biological system of interest. The downside of non-orthogonality is that components can be very similar to each other. As such, interpretation should be done with care: examining one component at a time can lead to wrong conclusions and plotting the loadings of two components against each other requires a transformation (`Supplementary Methods`). Possible additional constraints such as orthogonality and non-negativity of the loadings are described elsewhere [19, 20]. In this paper we will not constrain the PARAFAC models.

### Processing of longitudinal microbiome data

The main requirement of PARAFAC is that the residuals are distributed symmetrically around zero [15, 16, 19]. In this paper we perform the following four processing steps for each dataset to meet this requirement: centred log-ratio transformation, feature filtering, centring and scaling. In this section we outline these operations for multi-way data arrays (**Figure 2**). In the supplied R package, the function processDataCube() performs each of these steps according to user specifications.

**Figure 2.**
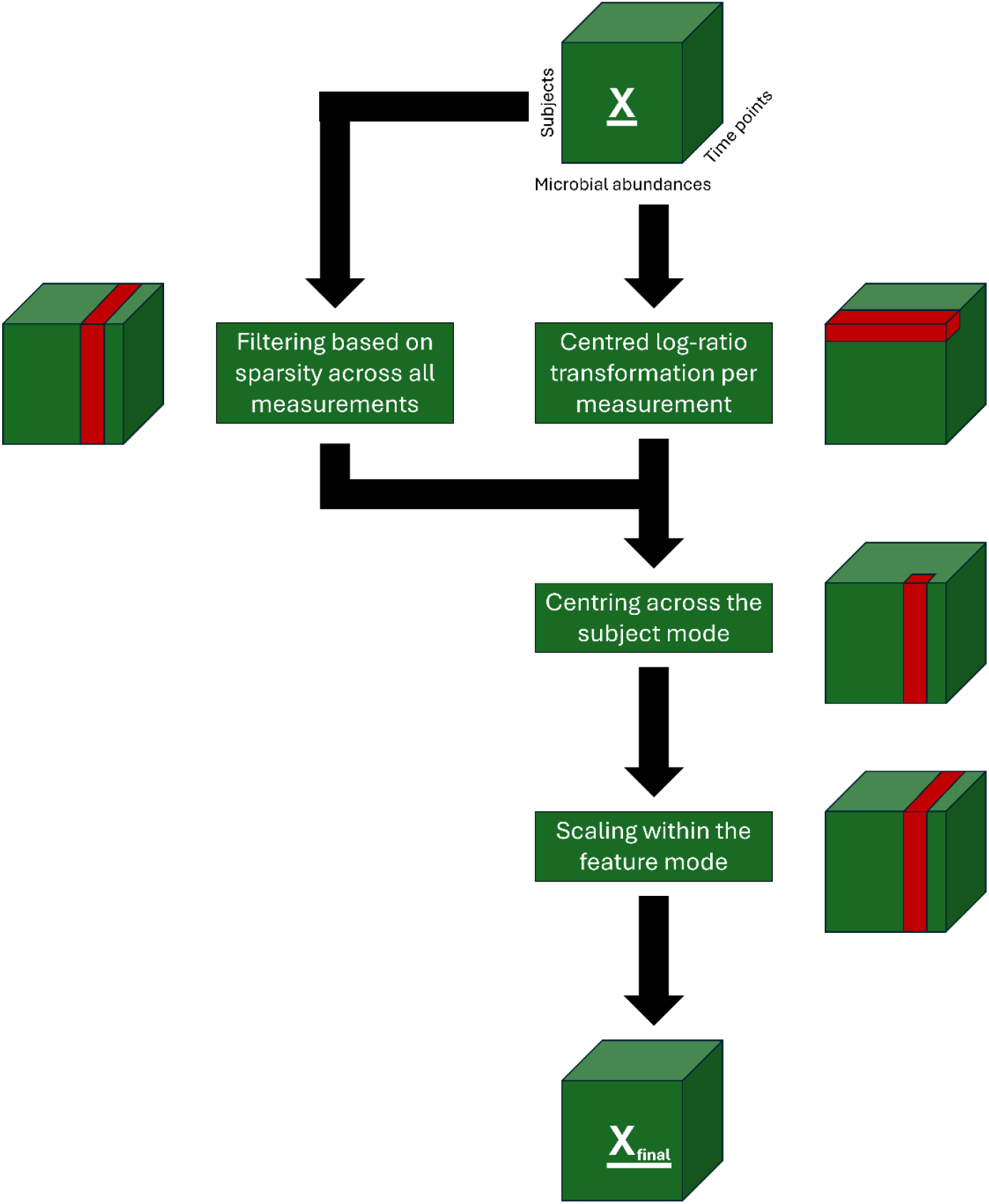
Workflow diagram of steps taken to process the microbiome count data for each dataset presented in this paper. The microbiome count data is indicated as <u>X</u> at the top of the figure. Per step, a small figure of the data cube is shown with the part of the cube that is used for the operation indicated in red. Sparsity-based feature filtering is based on the count data and is applied after the centred log-ratio transformation step. The final version of the data that is used as input for the PARAFAC modelling procedure is indicated as <u>X_final_</u> at the bottom of the figure.

Centred log-ratio transformation (CLR) is a popular method to create a version of the data that can be ordinated and interpreted through Euclidean distances [25]. This transformation approach deals with the compositionality inherent to microbiome data and makes the data continuous [26]. However, it is hampered by the zero-inflation commonly observed in microbiome data because zeroes are not defined in log space [27]. Here we choose to CLR transform the datasets after adding a pseudo-count of 1 to all counts (hereafter **<u>*X*^*clr*^</u>**), which appears to yield similar models as other pseudo-count strategies based on a brief sensitivity analysis (`Supplementary Figure 18`).

Due to the high dimensionality of the microbiome data, filtering of the features is a requirement to model the data appropriately and quickly [28, 29]. However, in longitudinal microbiome data the complete time trajectory of a feature should be considered prior to filtering. As such, we define the sparsity of a feature as the fraction of measurements where the feature has zero counts. A sparsity threshold can then be established to select features that are common, thus controlling the number of zeroes in the data. Sparsity-based feature filtering is based on the count data and is applied after the centred log-ratio transformation step. Since sparsity filtering is heavily dependent on the sequencing depth and microbiome complexity, no overall sparsity threshold can be suggested here. Therefore, a manual inspection of the sparsity per feature histogram is required to find a good balance between removing too many features and keeping features with enough variation of interest. The calculateSparsity() function in the supplied R package can be used for this purpose.

After CLR transformation and feature filtering, we used centring to remove offsets in the data [30, 31]. This can be done in three different directions for three-way arrays: across the subject mode, across the feature mode and across the time mode. Centring across the subject mode removes the average subject time profile from every combination of microbial abundance and time point, i.e.

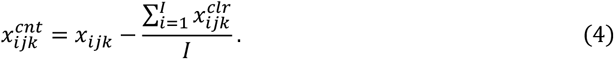

Centring across the feature mode removes the average feature time profile from every combination of subject and time point, i.e.

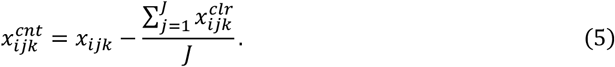

Centring across the time mode removes the average overall time profile from every combination of subject and microbial abundance, i.e.

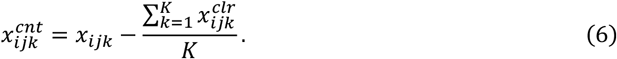

Here 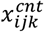 refers to the centred value of the *ijk*-th element in the data cube across the various directions specified above, and 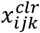 refers to the CLR transformed value of the *ijk*-th element. Both centring across the subject mode and across the feature mode affects the interpretation of the time mode as deviations from the average profile will be modelled. Since we expect individual offsets to be present in the exemplary studies and wish to focus on differences between the individuals, we centre across the subject mode. As a result, the PARAFAC models will describe deviations from the average time profile.

After centring, we scaled the data to change the importance attached to different parts [30, 31]. For three-way arrays, this involves scaling a two-way matrix to standard deviation one. As such, there are two possibilities for three-way arrays: scaling within the subject mode and scaling within the feature mode. Scaling within the subject mode makes the standard deviation for all counts across all subjects and microbial abundances per time point equal to one, i.e.

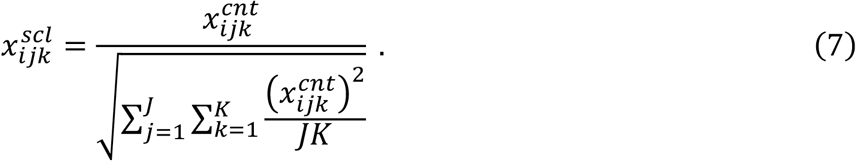

This makes all time points equally important for the modelling procedure. Scaling within the feature mode makes the standard deviation across all subjects and time points per microbial abundance equal to one, i.e.

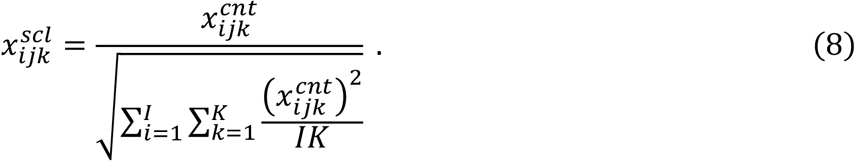

Here 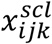 refers to the scaled value of the *ijk*-th element in the data cube within the various directions specified above and 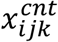 refers to the CLR transformed and centred value of the *ijk*-th element in the data cube. This makes all microbial abundances equally important for the modelling procedure. Since we are interested in groups of microbiota with different time profiles, we scale within the feature mode. The processing steps used for all datasets in this paper are outlined in **Figure 2**.

### Example dataset: Fujita et al., 2023

The first example is a long time series of experimental in vitro microbiomes [6]. Culture conditions, microbiome sampling, 16S rRNA gene amplicon sequencing and data pre-processing are described in the original publication. We analyse only one experimental set-up, the aquatic microbiome culture in a peptone medium, which was performed in eight replicates and measured daily for 110 days.

The amplicon sequence variant (ASV) data and taxonomic information were processed using the parafac4microbiome package (v1.0.3) in R (v4.4.1; [32]). The supplied dataset is a three-way array of counts of size 8 replicates x 28 microbial (ASV) abundances x 110 time points. Features were kept if they had ≤99% sparsity across the dataset (`Supplementary Figure 1`). This step resulted in 23 features being selected. We performed a centred log-ratio transformation with a pseudo-count of 1 to correct for compositionality [25, 26]. The data was then centred across the subject (here: replicate) mode and scaled within the feature mode to focus the analysis on describing differences from the average time trajectory. PARAFAC models of the processed data were created with parameters set to ctol=1e-6 and maxit=500. All models converged successfully by reaching the convergence tolerance threshold.

### Example dataset: Shao et al., 2019

The second example is a longitudinal faecal microbiome study with samples collected from 314 vaginally born and 282 caesarean-section born infants [7]. Sampling was performed at four time points post-partum: day 4, 7, 21, and a follow-up sample was taken between 4 to 12 months (referred to as the infancy period). Recruitment, microbiome sampling and 16S rRNA gene amplicon sequencing and pre-processing are described in the original publication.

Pre-processed sequencing data were obtained from ENA (accession numbers ERP115334 and ERP024601). Taxonomic classification was performed using mOTUs (v2.5.1; [33]). The marker-gene based operational taxonomic unit (mOTU) data and taxonomic information were processed using the `parafac4microbiome` package (v1.0.3) in R (v4.4.1; [32]). The supplied dataset is a three-way array of counts of size 395 subjects x 959 microbial (mOTU) abundances x 4 time points, keeping missing measurements as a row of NAs. This allows PARAFAC to impute the missing values. In total, 62.5% of the data was missing (`Supplementary Tables 1-2`). mOTUs were kept if the sparsity in either birth mode group was ≤90% (`Supplementary Figure 5`). This conserves biologically relevant mOTUs that only appear in one group and resulted in 91 of the 959 mOTUs being selected. The data was then centred log-ratio transformed using a pseudo-count of 1 to correct for compositionality [25, 26]. Subsequently, the data was centred across the subject mode and scaled within the feature mode to focus the analysis on describing differences from the average time trajectory. PARAFAC models of the processed data were created with parameters set to ctol=1e-8 and maxit=500 to ensure proper exploration of the loss space given the large amount of missing data. All models converged successfully by reaching the convergence tolerance threshold.

### Example dataset: Van der Ploeg et al., 2024

The third example is a longitudinal oral microbiome gingivitis intervention study [8]. Full details on the microbiome sampling and pre-processing are described in the original publication. We analyse only one oral microbiome niche, the supragingival plaque at the upper jaw lingual surfaces, which was measured in 41 subjects at 7 time points: day -14 (baseline), 0 (baseline, start of intervention), 2, 5, 9, 14 (end of intervention) and 21 (resolution). Using quantitative light-induced fluorescence [34–36], the subjects were stratified into low, medium and high responders based on the amount of red fluorescent plaque on day 14.

The ASV data and taxonomic information were processed using the `parafac4microbiome` package (v1.0.3) in R (v4.4.1; [32]). The supplied dataset is a three-way array of counts of size 41 subjects x 2253 microbial (ASV) abundances x 7 time points. In accordance with the original analysis, ASVs were kept if the sparsity was ≤50% in any response group. This step resulted in 65 of the 2253 ASVs being selected. Subsequently, a centred-log ratio transformation using a pseudo-count of 1 was performed to correct for compositionality [25, 26]. The data was then centred across the subject mode and scaled within the feature mode to focus the analysis on describing differences from the average time trajectory. PARAFAC models of the processed data were created with parameters set to ctol=1e-6 and maxit=500. All models converged successfully by reaching the convergence tolerance threshold.

### Selecting the appropriate number of PARAFAC components

Like in Principal Component Analysis, the number of biologically meaningful components that can be extracted from longitudinal microbiome data is limited. After a given number of components, PARAFAC will start to model noise. As such, one challenge of PARAFAC modelling for longitudinal microbiome data is selecting the appropriate number of components. A model with the correct number of components will (1) converge quickly, (2) not have highly similar loading vectors in any mode, (3) describe meaningful biological phenomena and (4) describe these phenomena even when one or more samples are removed from the data.

Various diagnostics and methods have been proposed and need to be manually examined to determine the appropriate number of components for a PARAFAC model [37]. We propose a two-step approach for selecting the correct number of components. First, we investigate quick convergence and similarity of the loading vectors by randomly initializing a number of PARAFAC models for 1-5 components. Next, we investigate the stability of the model using a jack-knife procedure while keeping the initialized vectors the same using a singular value decomposition. In the supplied R package, this procedure is automatically performed and visualized for inspection by the user with the functions assessModelQuality() and assessModelStability(), respectively. All output plots examined to select the number of components for each dataset are shown in `Supplementary Figures`.

### Metrics for assessing the appropriate number of components

For each dataset, we have investigated PARAFAC models with 1-5 components and a number of random initializations, depending on the size of the data and how quickly the models converge. In the supplied R package, this procedure is automatically performed and visualized for inspection by the user with the function assessModelQuality(). We examine the following metrics of interest.

In PCA, the residual sum-of-squares are plotted against the number of components in a scree-plot [38, 39]. The appropriate number of components is then the area of the plot where the residual sum-of-squares flattens out. For PARAFAC this principle holds as well [19].

The alternating least-squares implementation of PARAFAC that is used throughout this paper iteratively updates the loading vectors until the convergence threshold is reached. Therefore, the number of iterations needed to converge is a useful metric to inspect for the correct number of components. Randomly initialized PARAFAC models with the correct number of components should converge quickly compared to models with an extra component modelling noise [37, 40].

The core consistency diagnostic (CORCONDIA) indicates how well the assumed tri-linearity of the PARAFAC model holds [41]. The range of CORCONDIA is typically 0-100 but can be negative. The number of components in PARAFAC models with a CORCONDIA of 50-100 can be considered appropriate [19, 41].

The components of a PARAFAC model are non-orthogonal by default, which can sometimes lead to very similar loading vectors if the model cannot be estimated properly. The Tucker congruence coefficient identifies this similarity between pairs of loading vectors of the same mode [42, 43]. A value in the range of 0.85-0.94 suggests that the loading vectors are fairly similar, whereas a value of 0.95-1 means that the loading vectors are essentially equal [42]. The Tucker congruence coefficient in the `parafac4microbiome` package uses the implementation from the multiway R package (version 1.0-6; [20]).

### Assessing model stability through the jack-knifing of samples

We expect a PARAFAC model with the correct number of components to describe meaningful biological phenomena, even when one or several samples are removed from the data. We test this by performing iterative jack-knifing of samples [44]. In the supplied R package, this procedure is automatically performed and visualized for inspection by the user with the function assessModelStability(). With this jack-knifing approach, one random sample is removed in every iteration, after which a PARAFAC model is fitted to the data. We perform balanced jack-knifing, removing one sample from every subject group in every iteration, if such subject metadata is available. If this is impossible due to unequally sized groups, samples are randomly chosen. We choose to initialise the modelling algorithm using a singular value decomposition for every iteration to ensure that instability can only be caused by the removed sample(s). A PARAFAC model with the correct number of components will be stable (i.e. have a small standard deviation for every loading) across all jack-knifing iterations.

### Post-hoc clustering of PARAFAC loadings

Clustering of the feature loadings can help identify groups of microbiota that are connected to a particular time profile in the PARAFAC model. We performed the clustering procedure of the feature loadings as follows. In accordance with the original analysis in van der Ploeg et al. [8], features were not considered if the variation explained by the model for their entire time trajectory was lower than the average or the congruence loading of the feature was lower than 0.4 [45]. The remaining features were then clustered based on their modelled time trajectories using the K-medoids algorithm from the cluster R package (v2.1.4; [46]) with 50 random starts to be robust against outliers. The number of clusters was determined using the within-cluster sum of squares, silhouette width and gap statistic metrics as reported by the factoextra R package (`Supplementary Figure 17`; v1.0.7; [47]).

## Results

### PARAFAC can identify the main time-resolved variation in longitudinal microbiome data

The first example is a long time series of experimental in vitro microbiomes [6]. We analyse only one experimental set-up, the aquatic microbiome culture in a peptone medium, which was performed in eight replicates and measured daily for 110 days. With this example, we aim to visualize the main time-resolved variation that can be modelled through PARAFAC.

We inspected the unprocessed count data by computing the relative abundances for every replicate culture at every time point (**Figure 3A**). In most samples, 2 amplicon sequence variants (ASVs) belonging to *Aeromonas* spp. comprised most of the counts. The exceptions were: the emergence of *Cedecea, Burkholderia-Caballeronia-Paraburkholderia*, and *Pandoraea* species between days 50-110 in replicate culture 4, the emergence of *Clostridioides mangenotii* and a *Lachnoclostridium* species between days 80-100 in replicate cultures 4 and 8, the disappearance of a *Salmonella, Propionivibrio, Microbacter* and *Herbaspirillum* species between days 0-60 in replicate culture 3, the transient appearance of a *Microbacter* species in replicate cultures 2 and 8, and a (small) increase in a *Salmonella* species between days 100-110 in most replicate cultures. There was no clear time profile visible that all replicates had in common.

**Figure 3.**
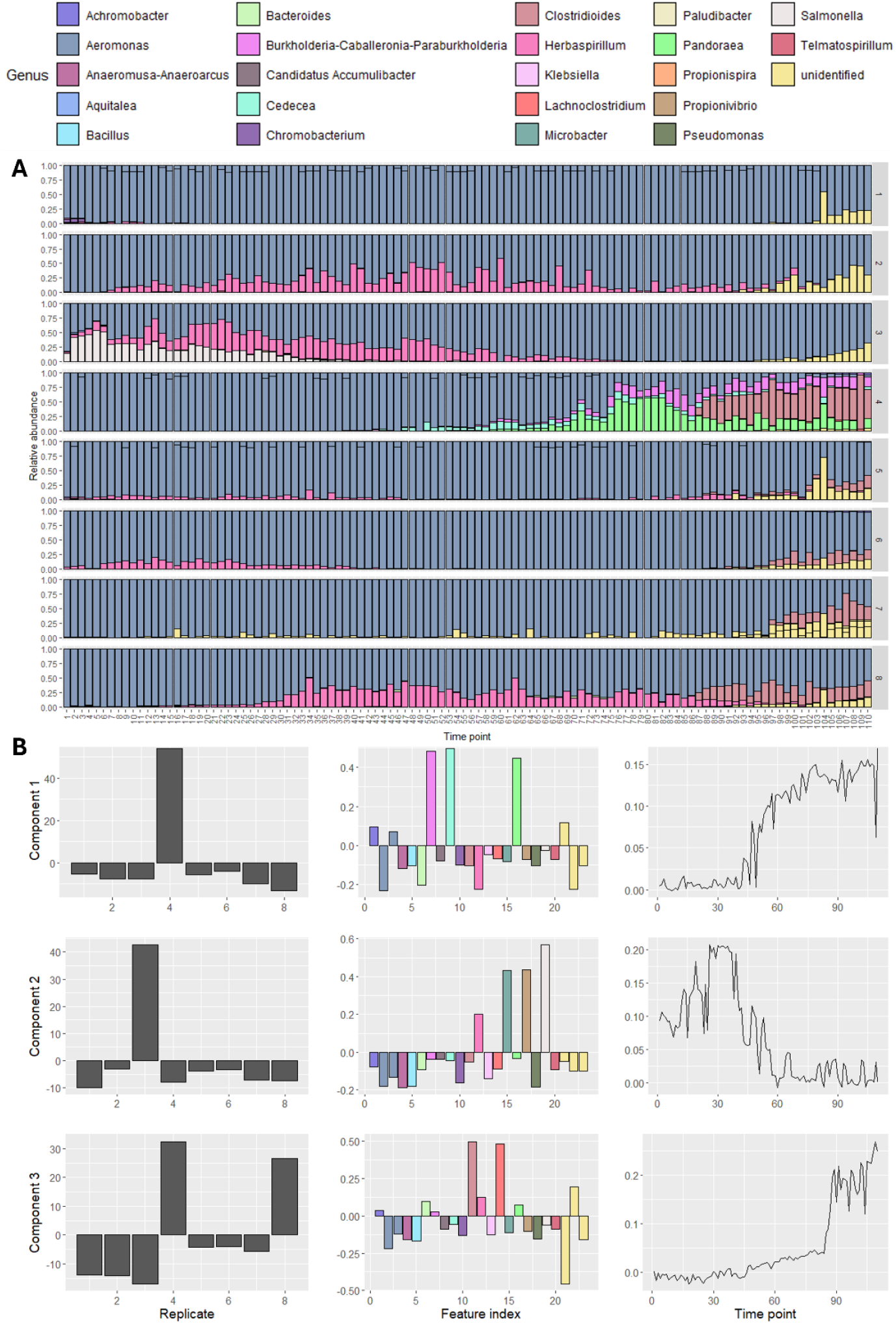
Overview of the Fujita et al., 2023 dataset. (A) Relative abundance plot of the dataset, split per replicate. In most samples, Aeromonas species comprise the vast majority of the counts. In the latter half of the samples in replicate 4 and the first half of the samples in replicate 3 other species are present that comprise the majority. There is no clear time profile visible that all replicates have in common. (B) The best three-component PARAFAC model of the processed data describing 39.0% of the variation. The components are shown in the rows, while the replicate, microbial abundance and time modes are shown in the columns. The first component described the emergence of a Cedecea, Burkholderia-Caballeronia-Paraburkholderia and Pandoraea species between days 50-110, which is unique to replicate 4. The second component described the disappearance of a Salmonella, Propionivibrio, Microbacter and Herbaspirillum species between days 0-60, which is unique to replicate 3. The third component described the emergence of Clostridioides mangenotii and a Lachnoclostridium species between days 80-100 in replicates 4 and 8.

We inspected PARAFAC modelling performance and stability for the processed count data using 1-5 components. The core consistency diagnostic (CORCONDIA) scores dropped below 80 for 5 components. The three and four-component models explained 38.78% and 45.08% on average, respectively (`Supplementary Figure 2`). Performing a jack-knifing procedure showed that the three-component PARAFAC models were more stable than the four-component models (`Supplementary Figures 3-4`). Hence, we selected the model explaining the most variation from all randomly initialized models of three components. The selected model explained 39.0% of the variation (**Figure 3B**).

As expected, the selected PARAFAC model did not describe *Aeromonas* ASVs, because these do not vary strongly over time. The first component described the emergence of a *Cedecea, Burkholderia-Caballeronia-Paraburkholderia* and *Pandoraea* species between days 50-110, which is unique to replicate 4. The second component described the disappearance of a *Salmonella, Propionivibrio, Microbacter* and *Herbaspirillum* species between days 0-60, which is unique to replicate 3. The third component described the emergence of *Clostridioides mangenotii* and a *Lachnoclostridium* species between days 80-100 in replicates 4 and 8. Some of these microbiota may have low relative abundances but can still describe meaningful differences between the replicates.

While this example highlights the strength of PARAFAC: identifying and modelling the largest sources of variation in a longitudinal microbiome dataset, the PARAFAC model only described the variation partially (39%). As a result, the transient appearance of a *Herbaspirillum* species in replicate cultures 2, 3 and 8, nor the (small) increase in a *Salmonella* species between days 100-110 in most replicate cultures, were not modelled.

### PARAFAC finds differences between subject groups and enhances comparative analyses

The second example is a longitudinal infant faecal microbiome study [7]. Samples were collected from 314 vaginally born and 282 caesarean-section born infants at four time points: 4, 7 and 21 days after birth and a follow-up sample during infancy (between 4-12 months after birth). One of the four measurements is missing for most infants in this cohort, and we keep the missing data in the data cube for the PARAFAC algorithm to impute (`Supplementary Table 1-2`). With this example, we show how PARAFAC can identify the differences between the subject groups and enhance the comparative analysis done by the original authors despite a moderate amount of missing data.

The authors stated that there was a very clear distinction in the microbiomes of vaginally and caesarean-section born children, also over time. To analyse this distinction, we inspected the unprocessed count data by computing the average relative abundance for the vaginal and caesarean groups at every time point (**Figure 4A**). This confirmed a clear difference at the phylum level between the vaginal and caesarean groups at day 4, 7 and 21 postpartum, which disappeared in infancy.

**Figure 4.**
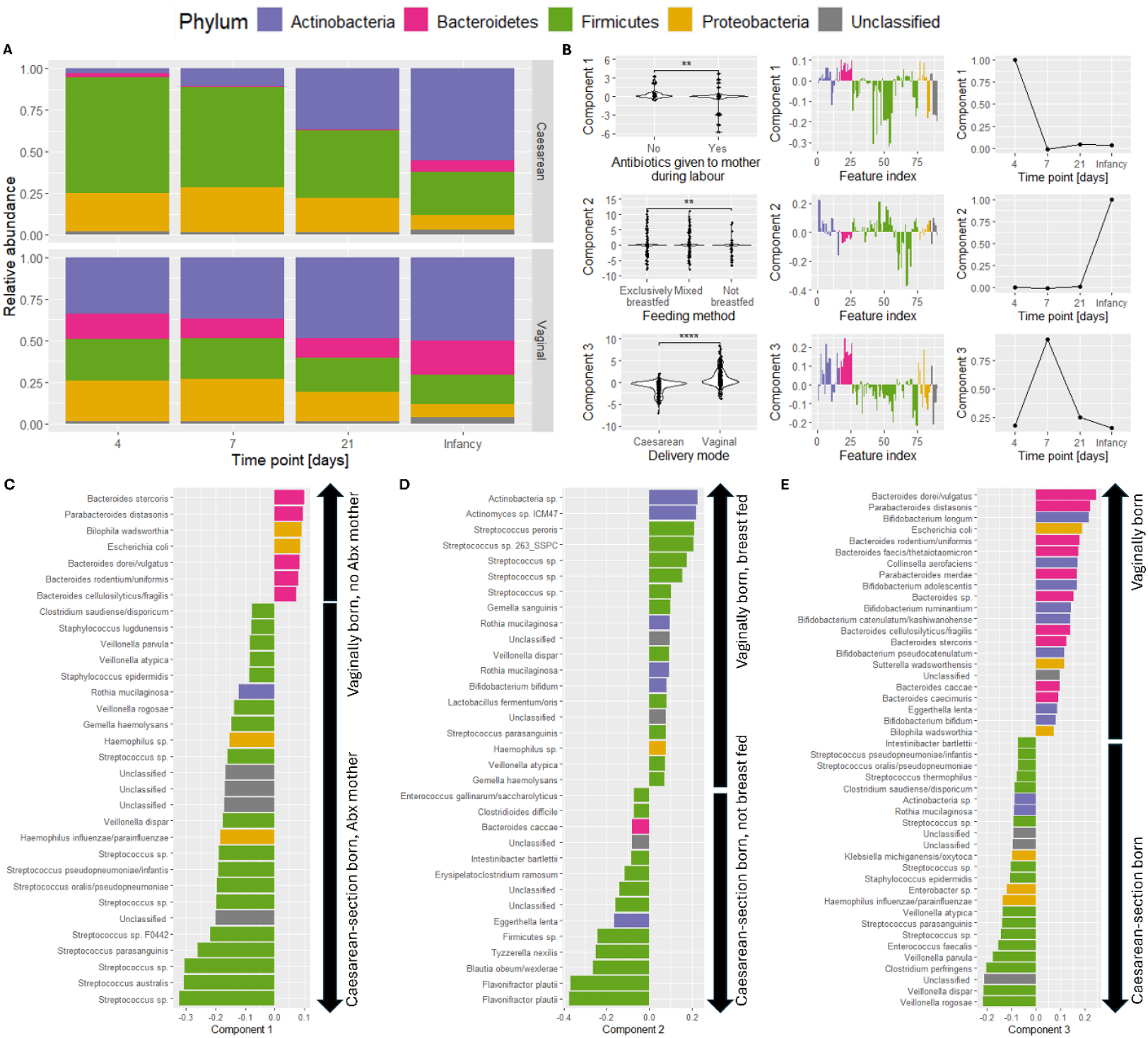
Overview of the Shao et al., 2019 dataset. (A) Overview of the average relative abundance per time point in vaginal and caesarean-section born infants. The birth mode groups are distinguishable between time points 4 and 21 postpartum and become similar at infancy. (B) The best three-component PARAFAC model of the processed data, explaining 9.40% of the variation. Components 1-3 all describe differences between the birth mode groups. Component 1, mainly present on day 4, additionally describes differences based on antibiotics being given to the mother during labour. Component 2, mainly present during infancy, additionally describes feeding mode. Component 3 describes only describes differences between the birth mode groups, which are present throughout the study but are largest on day 7. Wilcoxon rank-sum tests were used to compare the subject loadings against subject metadata. Full results of the tests are supplied in Supplementary Table 3. (C) Loadings of the microbiota in component 1. Positively loaded microbiota are associated with vaginally born infants and/or mothers having received antibiotics during labour. Negatively loaded microbiota are associated with caesarean-section born infants and/or mothers not having received antibiotics during labour. (D) Loadings of the microbiota in component 2. Positively loaded microbiota are associated with vaginally born and/or breast-fed infants. Negatively loaded microbiota are associated with caesarean-section born and/or not breast-fed infants. (E) Loadings of the microbiota in component 3. Positively loaded microbiota are associated with vaginally born infants. Negatively loaded microbiota are associated with caesarean-section born infants. Feature loadings in (C-E) are filtered to only show loadings with an absolute value higher than 0.075. For the unfiltered version of these plots, see Supplementary Figures 9-11.

We inspected PARAFAC modelling performance and stability for the processed count data using 1-5 components. The CORCONDIA scores dropped below 80 at 3-5 components (`Supplementary Figure 6`). Performing a jack-knifing procedure showed that the two-component model was as stable as the three-component model (`Supplementary figures 7-8`). Hence, we selected the model explaining the most variation from all randomly initialized models of three components. The selected model explained 9.40% of the variation (**Figure 4B**).

All components of the selected model described differences in birth mode (p=2.4e-4, p=7.3e-9 and p=1.2e-46, respectively; Benjamini-Hochberg corrected Wilcoxon rank-sum test; `Supplementary Table 3`) and differences based on the *Bacteroides* profile as described by the original authors based on the median relative abundance of the genus (p=1.5e-4, p=7.3e-9, p=9.2e-59, respectively; [7]). In addition, the first component described differences related to whether antibiotics were given to the mother during labour (p=0.0059) and the second component described differences based on feeding method (p=0.024).

Inspection of the positive loadings in the first component revealed that PARAFAC described the genera that were enriched in vaginally born infants: *Bacteroides stercoris, Parabacteroides distasonis, Bilophilia wadsworthia, Escherichia coli, Bacteroides dorei/vulgatus, Bacteroides rodentium/uniformis and Bacteroides cellulosilyticus/fragilis* (**Figure 4C**). These genera were depleted in caesarean-section born infants and infants whose mothers received antibiotics during labour. The first component also described genera that were enriched in caesarean-section born infants or infants whose mothers received antibiotics during labour: *Streptococcus australis, Streptococcus parasanguinis, Streptococcus oralis*, amongst others. These genera were depleted in vaginally born infants.

Inspection of the positive loadings in the second component revealed that PARAFAC described the genera that were enriched in vaginally born or breast-fed infants: *Actinobacteria* sp., *Actinomyces* sp. ICM47, and *Streptococcus peroris*, amongst others (**Figure 4D**). These genera were depleted in caesarean-section born and formula-fed infants. It also described the genera that were enriched in caesarean-section born or formula-fed infants: *Flavonifractor plautii, Blautia obeum/wexlerae* and *Tyzzerella nexilis*, amongst others. These genera were depleted in vaginally born and breast-fed infants. While not described in the original analysis, these results partially agree with other literature [46–48].

Inspection of the positive loadings in the third component revealed that PARAFAC described the genera that were enriched in vaginally born infants: *Bacteroides dorei/vulgatus, Parabacteroides distasonis* and *Bifidobacterium longum*, amongst others (**Figure 4E**). These genera were depleted in caesarean-section born infants. It also described the genera that were enriched in caesarean-section born infants: *Veillonella rogosae, Veillonella dispar*, and *Clostridium perfringens*, amongst others. These genera were depleted in vaginally born infants. This result is consistent with the original analysis and other studies investigating the development of the infant gut microbiome in hospital environments [48, 49].

Taken together, the PARAFAC model of this example dataset confirmed previously found differences between the caesarean-section and vaginally born infants. It also found novel genera that describe the differences between the birth mode groups and feeding mode groups. Additionally, the moderate (62.5%) amount of missing data did not hamper the modelling procedure. In summary, the PARAFAC model of this dataset confirms and enhances the results from the comparative analysis done by the original authors.

#### Post-hoc clustering of PARAFAC results can help identify microbial groups of interest

The third example is a longitudinal oral microbiome gingivitis intervention study [8]. Here, we analyse one oral microbiome niche, the supragingival plaque at the upper jaw lingual surfaces, which was measured in 41 subjects at 7 time points: day -14 (baseline), 0 (baseline, start of intervention), 2, 5, 9, 14 (end of intervention) and 21 (resolution). Using quantitative light-induced fluorescence [34–36], the subjects were stratified into low, medium and high responders based on the amount of red fluorescent dental plaque on day 14. With this example, we show how a post-hoc clustering approach of a PARAFAC model can help identify groups of microbiota related to the intervention.

We inspected PARAFAC modelling performance and stability for the processed count data using 1-5 components. The CORCONDIA scores dropped below 80 for 5 components and the sum of squared errors began to flatten between 2-3 components (`Supplementary Figure 13`). The three-component models had very similar feature and time mode loadings between several components, making them not suitable for describing the data (`Supplementary Figure 14`). This was confirmed by the jack-knifing procedure (`Supplementary Figures 15-16`). Hence, we selected the model explaining the most variation from all randomly initialized models of two components. The selected model explained 19.21% of the variation (**Figure 5A**).

**Figure 5.**
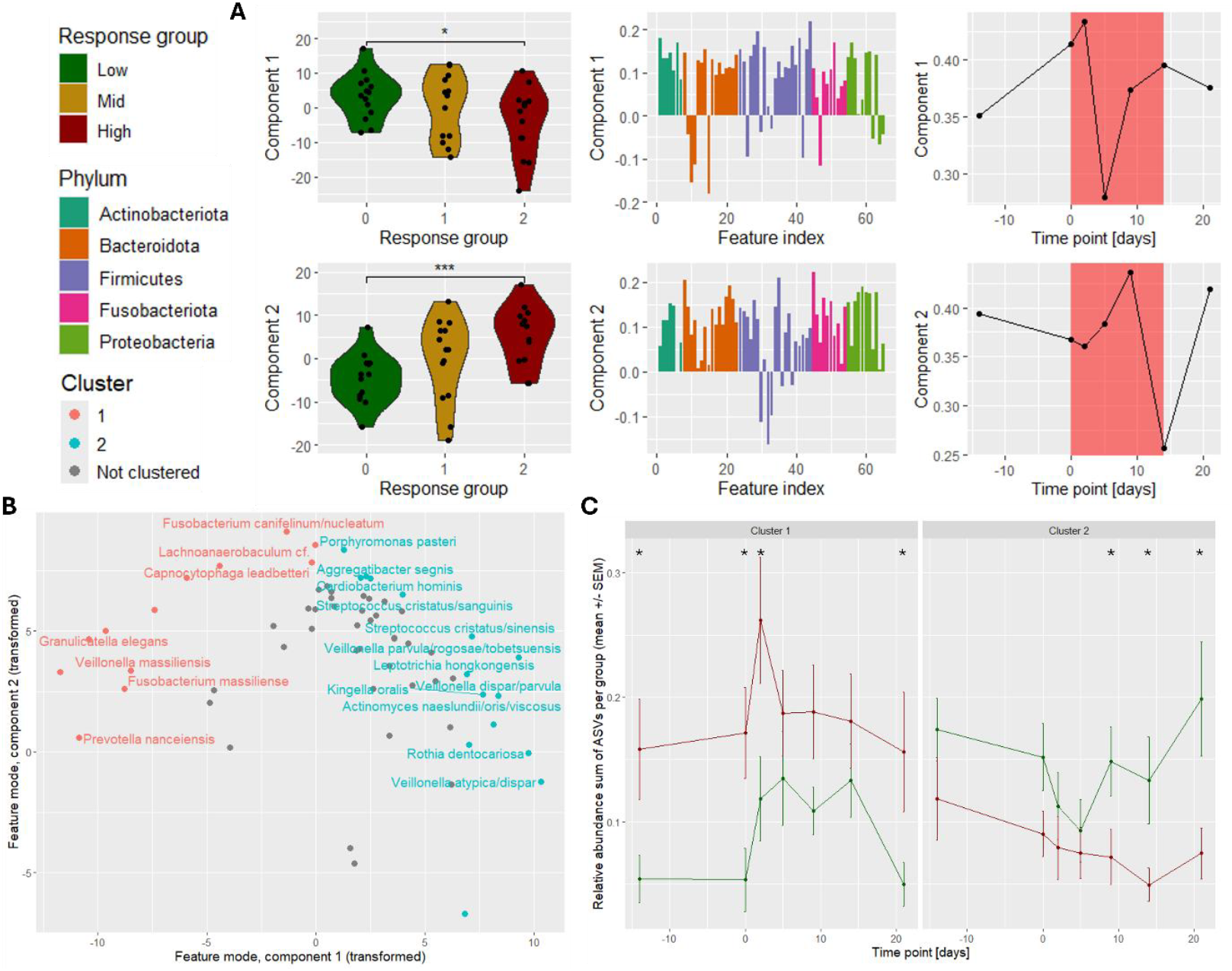
Overview of the van der Ploeg et al., 2024 dataset. (A) The best two-component PARAFAC model of the processed data, explaining 19.21% of the variation. Both components describe differences between the response groups that are relatively constant throughout the study. The time profile in component 1 describes that the differences between the response groups are larger on day 2 and much smaller on day 5. The time profile in component 2 describes that the differences between the response groups are larger on day 9 and much smaller on day 14. Wilcoxon rank-sum tests were used to compare the subject loadings against subject metadata. Full results of the tests are supplied in Supplementary Table 4. (B) Scatter plot showing the transformed feature loadings (Supplementary Methods) of PARAFAC component 1 versus component 2. Features were clustered into two groups after filtering. Only amplicon sequence variants (ASVs) with taxonomic annotations resolved to species level are labelled. (C) Overview of the relative abundance sum per ASV cluster in the low and high response groups. Error bars correspond to the standard error of the mean (SEM) across all samples per response group. The mean difference between the high and low response groups per time point was tested using a Benjamini-Hochberg corrected permutation test of 999 iterations (*: p≤0.05; **: p≤0.01; ***: p≤0.001). Mid responders are not shown for visual clarity.

To interpret the models, we tested the correlation of the subject loadings with the measured parameters of gingivitis. This approach revealed that both components of the PARAFAC model significantly described the amount of red fluorescent plaque over time (p=1.8e-5 and p=0.029, respectively; Benjamini-Hochberg corrected Pearson correlation test; `Supplementary Table 4`) and the percentage of sites that were covered by plaque (p=0.029 and p=0.029, respectively).

Following the original analysis [8], we identified microbial subcommunities with common responses by selecting and then clustering only well-modelled amplicon sequence variants (ASVs) based on their fitted response using a K-medoids algorithm (**Figure 5B**). The sum of the relative abundances of the ASVs in each cluster was then determined to test the difference between the microbiomes of individuals in the low and high response groups at every time point. This approach revealed that the relative abundance sum of ASV cluster 1 was significantly higher in the high responder individuals compared to the low responder individuals on days -14, 0, 2, and 21 (**Figure 5C, Supplementary Table 5**; p≤0.05; Benjamini-Hochberg corrected permutation test of mean difference). Similarly, the relative abundance sum of ASV cluster 2 was found to be significantly higher in low responder individuals compared to high responder individuals on days 9, 14 and 21. Taken together, the PARAFAC models of plaque microbiomes described microbial subcommunities with common dynamics that connect to the magnitude of the gingivitis response.

## Discussion

Using three example studies [6–8], we show that the unsupervised modelling approach of Parallel Factor Analysis (PARAFAC) can (1) identify the main time-resolved variation in longitudinal microbiome data, (2) find differences between subject groups and enhance comparative analyses and (3) helps identify feature groups of interest using a post-hoc clustering approach. In addition, we show that PARAFAC is robust to missing measurements in the Shao et al., 2019 [7] example. Robustness of the PARAFAC algorithm in the context of missing data was an expected result and is well-documented [50, 51]. We also show in the Shao et al., 2019 [7] and van der Ploeg et al., 2024 [8] examples that the modelled variation is related to clinically relevant parameters. However, many other multi-way approaches exist that are more flexible in their modelling procedure [31]. For example, Tucker3 allows for interactions between the components [52]. PARAFAC2 allows for time shifts that are unique per subject [19, 53]. Multi-way regression, in which the PARAFAC components are supervised using a response variable, is also possible with N-way partial least squares [54]. As highlighted in the Fujita et al. example study [6], these multi-way dimension reduction methods require commonality between the subjects to be useful. While PARAFAC’s treatment of time as a categorical variable is useful to describe time profiles in small-scale microbiome studies, large-scale microbiome studies should be analysed with models that use quantitative time factors, such as General Additive Models [55] or (spline) Linear Mixed Effects Models [56, 57]. Further research is needed to investigate the relevance of these methods in the context of longitudinal microbiome data analysis.

In PARAFAC, the nature of the third mode in the multi-way array is not limited to time. For example, a recent study has shown that soil treatment can be used as the third mode to identify relationships between the abundances of mycorrhizal fungi and plant diversity and history [58]. Additionally, the principles shown in this work are generalizable to four or more modes if that better represents the data that was generated by the study design.

In this work we have not elaborated on the various options that exist for processing microbial abundance data. The sparsity filtering approach presented in this work does not consider microbiota disappearing as time progresses. The addition of a pseudo-count prior to a centred log-ratio (CLR) transformation is a complex and unresolved matter [59, 60]. It has been noted that this approach introduces a dependency with library size, depending on the number of zeroes in the data [59]. Other work [60] suggests that a sample-specific pseudo-count may resolve this issue, but further research is needed to examine this approach in the context of longitudinal microbiome data. Other options include setting a pseudo-count value of 0.5, 0.1 or replacing the zeroes with a random value between 0 and 1 drawn from a uniform distribution. We conducted a brief sensitivity analysis for these options (`Supplementary Figure S18`) and decided to use a pseudo-count of 1 as this treats all zeroes as relevant values while yielding similar models. In a related paper on tensor factorization of longitudinal microbiome data [61] the robust-centred log-ratio was suggested as an approach to transform the data and to correct for compositionality. This approach instead calculates the geometric mean by ignoring the zeroes and setting them to missing values after the transformation. This makes all zeroes available for imputation by the PARAFAC model, but ignores the different nature that zeroes can potentially have [27]. Further research is needed to investigate the appropriateness of the various processing methods for (longitudinal) microbiome count data in describing biological phenomena of interest.

## Conclusions

We have shown that Parallel Factor Analysis is an applicable method for longitudinal microbiome data analysis across a wide range of microbial environments. Our results demonstrate that PARAFAC is able to (1) model the main time-resolved variation in longitudinal microbiome data,

1. find differences between subject groups and enhance comparative analyses despite a moderate amount of missing data, and (3) help identify feature groups of interest. The analyses and the example datasets with the resulting figures are implemented in the R package `parafac4microbiome`, which is available on CRAN at https://cran.rstudio.com/web/packages/parafac4microbiome/.

## List of abbreviations

ALS: Alternating Least-Squares
ASV: Amplicon Sequence Variant
CLR: Centred Log-Ratio
CORCONDIA: Core Consistency Diagnostic
mOTU: Marker-gene based Operational Taxonomic Unit
PCA: Principal Component Analysis
PARAFAC: Parallel Factor Analysis

## Ethics approval and consent to participate

Not applicable.

## Consent for publication

Not applicable.

## Availability of the data and materials

The package used during the current study is available in the GitHub repository, https://doi.org/10.5281/zenodo.13862690. The datasets generated and/or analysed during the current study are available in the GitHub repository, https://doi.org/10.5281/zenodo.11072651. A STORMS (Strengthening The Organizing and Reporting of Microbiome Studies)[62] checklist is available at https://doi.org/10.5281/zenodo.11072651.

## Authors*’* contributions

G.R. van der Ploeg: conceptualization, visualization, formal analysis, methodology, programming, writing – original draft; J.A. Westerhuis: conceptualization, methodology, supervision, formal analysis, writing – original draft; A. Heintz-Buschart: conceptualization, formal analysis, supervision, project administration, writing – original draft; A.K. Smilde: conceptualization, methodology, supervision, funding acquisition, writing – review & editing.

## Acknowledgements

We want to thank Jesse Alderliesten (UU/UvA) for their useful discussions and suggestions.

