## Supplementary Information for "parafac4microbiome: Exploratory analysis of longitudinal microbiome data using Parallel Factor Analysis"

### Supplementary Methods

*Transforming PARAFAC loadings for scatter plots*

Due to non-orthogonality, scatter plots of PARAFAC loadings require a transformation to find an orthonormal basis [1]. To create a scatter plot of the subject loadings in the matrix $\boldsymbol{A}$, we compute

$$\widetilde{\boldsymbol{A}} = \boldsymbol{A}(\boldsymbol{T}')^{-1}$$

where $\boldsymbol{T} = \boldsymbol{F}^{+}\widetilde{\boldsymbol{F}}$, $\boldsymbol{F} = \boldsymbol{C} \odot \boldsymbol{B}$ and $\odot$ denotes the column-wise Kronecker (or Khatri-Rao) product [2], $\boldsymbol{F}^{+}$ is the pseudoinverse of $\boldsymbol{F}$, and $\widetilde{\boldsymbol{F}}$ is an orthonormalized version of $\boldsymbol{F}$ through Gram-Schmidt orthonormalization [3]. Analogously, the feature loadings in the matrix $\boldsymbol{B}$ can be plotted after computing

$$\widetilde{\boldsymbol{B}} = \boldsymbol{B}(\boldsymbol{T}')^{-1}$$

where $\boldsymbol{F} = \boldsymbol{A} \odot \boldsymbol{C}$ and for the time loadings in the matrix C

$$\widetilde{\boldsymbol{C}} = \boldsymbol{C}(\boldsymbol{T}')^{-1}$$

where $\boldsymbol{F} = \boldsymbol{B} \odot \boldsymbol{A}$. After the transformation, loadings from two components can be plotted against each other like in PCA. In this paper we used the pracma R package package (version 2.4.2; [4]) for pseudo-inverse calculation and Gram-Schmidt orthonormalization. This transformation functionality is available in our R package with the function transformPARAFACloadings().

### Supplementary Figures

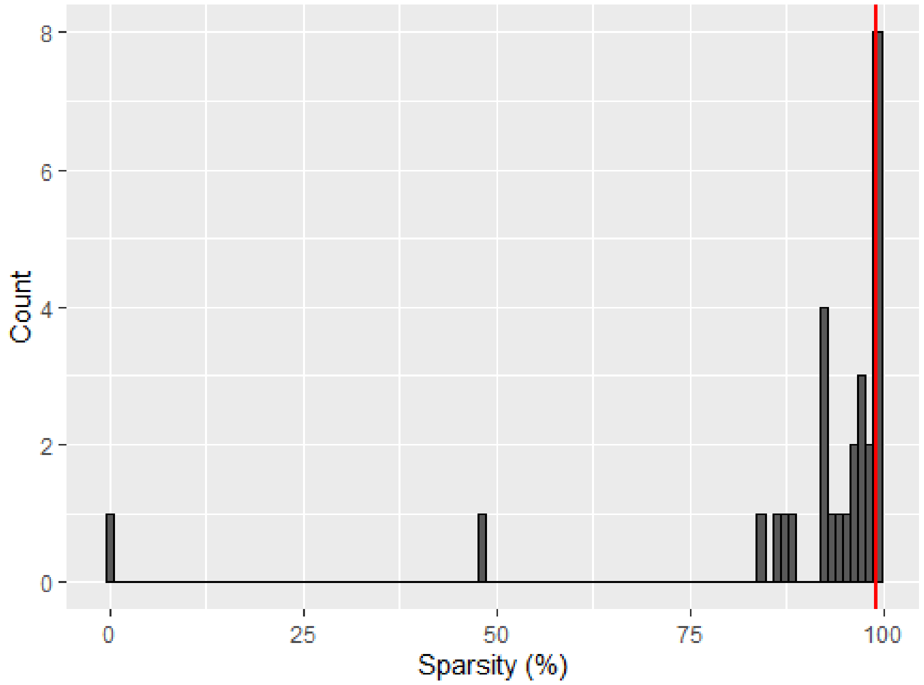


*Supplementary Figure 1: histogram displaying the sparsity of the unprocessed microbial abundance count data in the Fujita et al., 2023 dataset. Sparsity is defined as the percentage of measurements with a zero count. Microbial abundances were used for modelling if they have a sparsity ≤99%, indicated by the vertical red line. 28 microbial abundances were checked this way and 23 remained after this filtering step.*

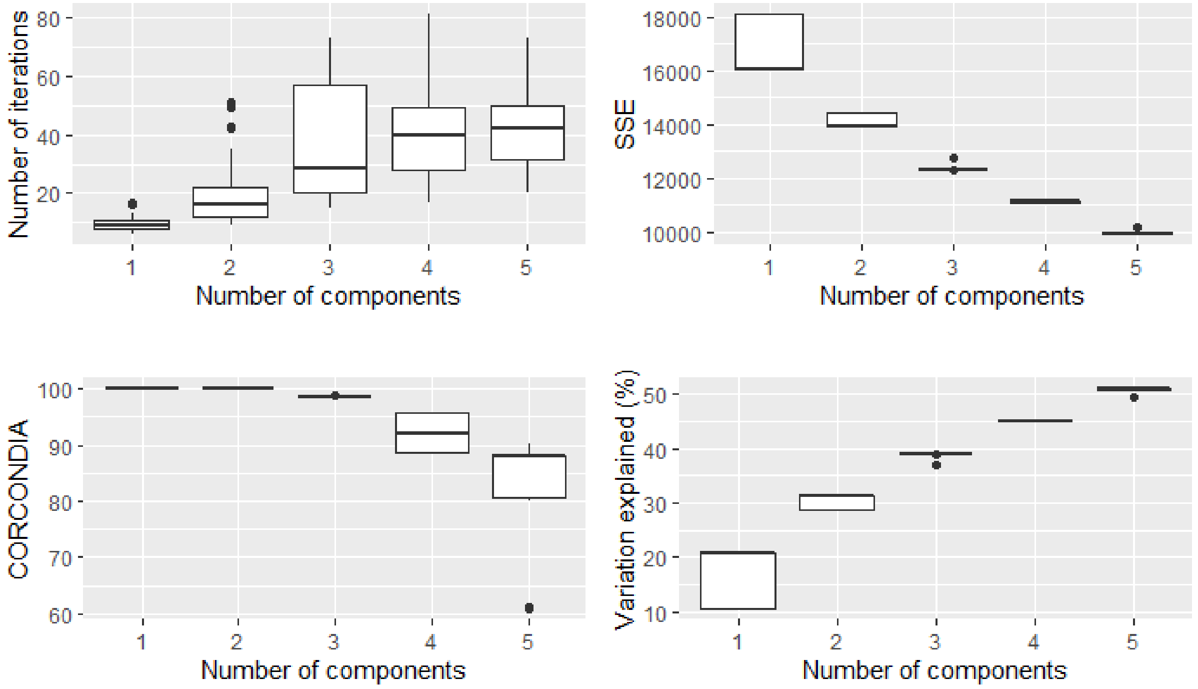


*Supplementary Figure 2: overview plots showing the number of iterations needed, sum of squared error, CORCONDIA score and variance explained for 50 randomly initialized models of 1-5 components each of the processed Fujita et al., 2023 dataset. The CORCONDIA scores dropped below 80 at 5 components.*

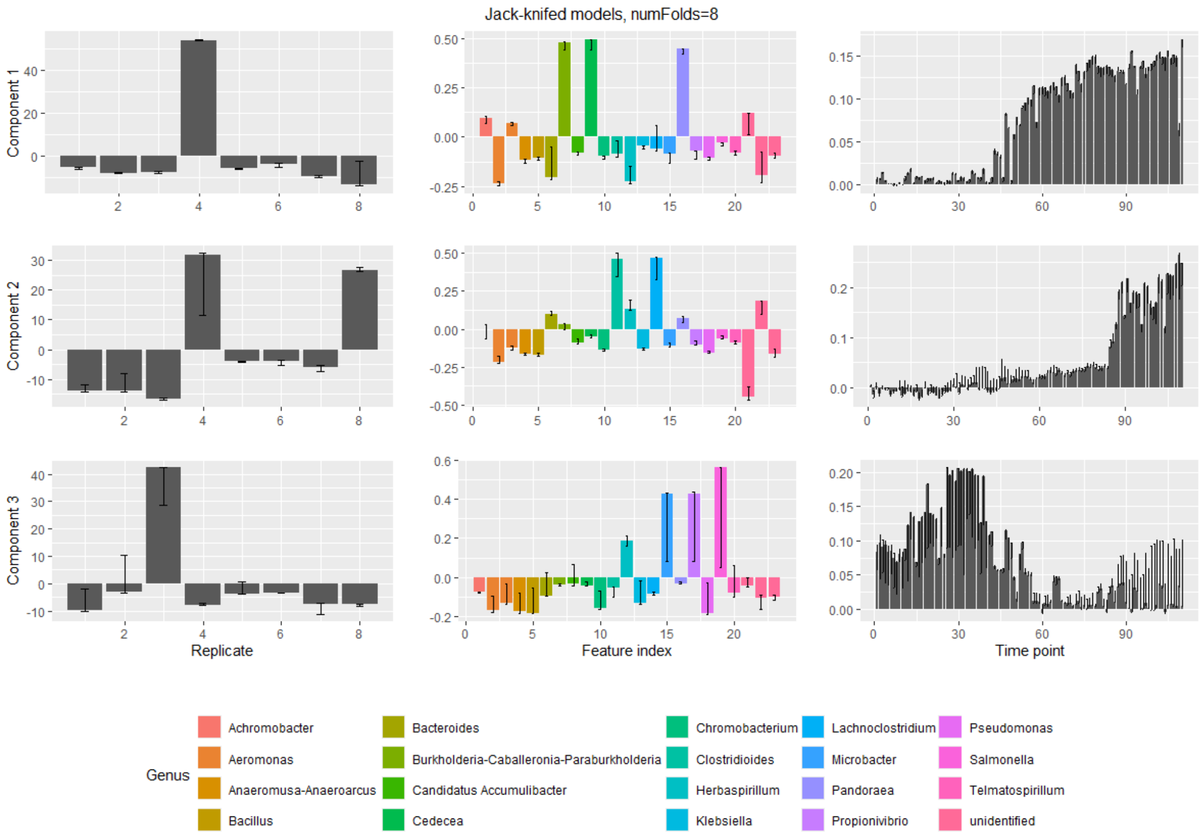


*Supplementary Figure 3: summary of the 8 jack-knifed models of three components created as part of the jack-knifing procedure for the processed Fujita et al., 2023 dataset. Modes are shown in the columns (from left to right: replicate mode, feature mode, time mode) and components are shown in the rows. Error bars display the range of all values seen for a particular loading across all initialized models. This number of components resulted in stable loadings across all modes.*

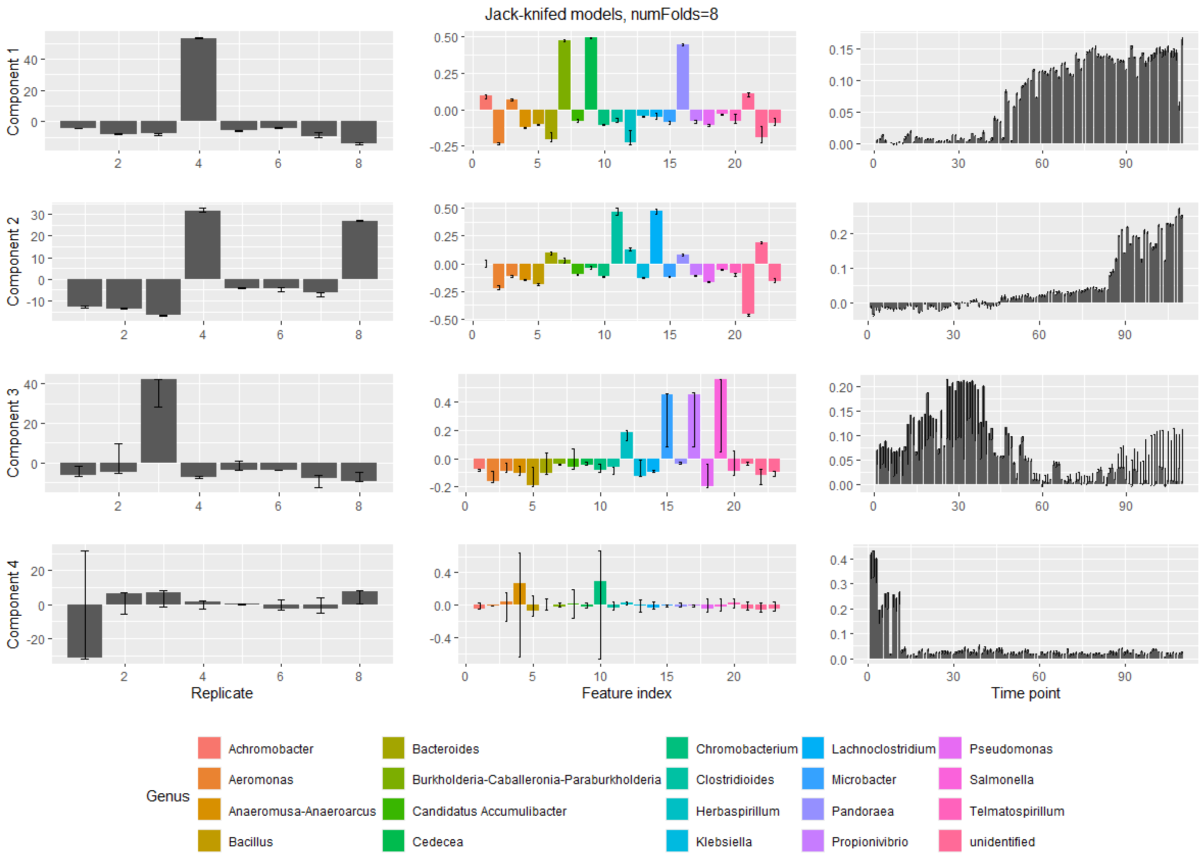


*Supplementary Figure 4: summary of the 8 jack-knifed models of four components created as part of the jack-knifing procedure for the processed Fujita et al., 2023 dataset. Modes are shown in the columns (from left to right: replicate mode, feature mode, time mode) and components are shown in the rows. Error bars display the range of all values seen for a particular loading across all initialized models. These models show instability in component 4, mainly in the subject and feature mode.*

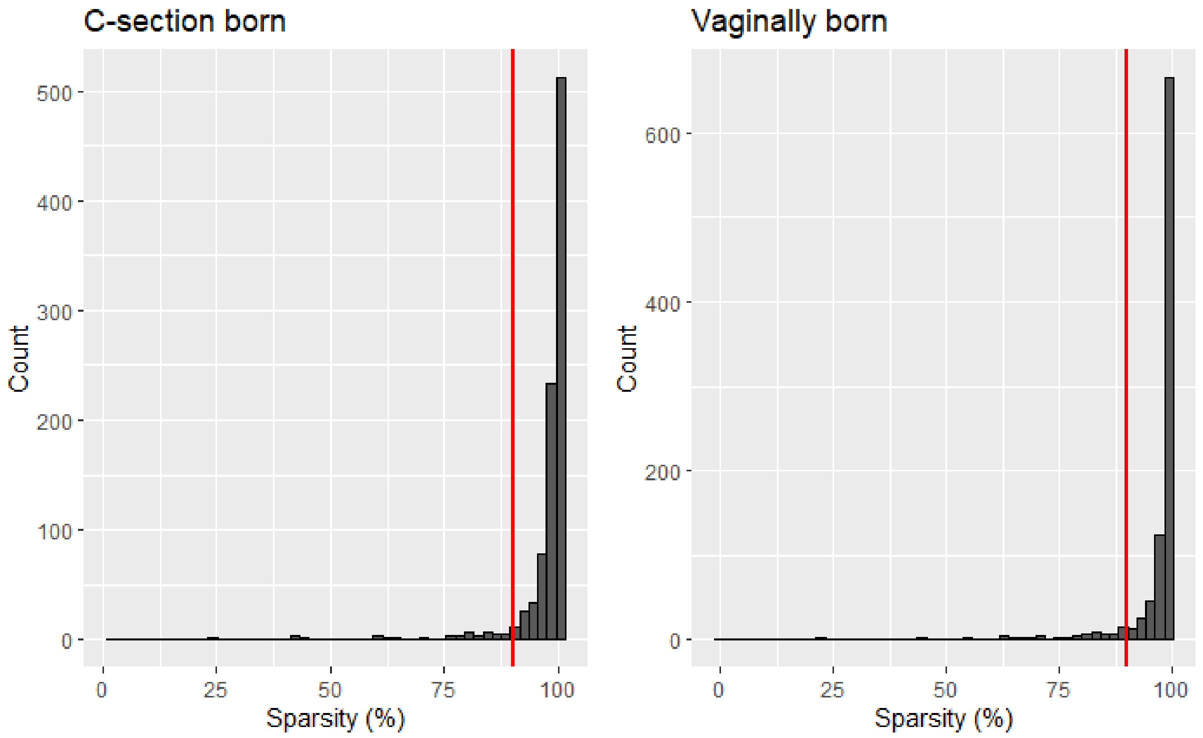


*Supplementary Figure 5: histogram displaying the sparsity of the unprocessed microbial abundance count data in each birth mode group of the Shao et al., 2019 dataset. Sparsity is defined as the percentage of measurements with a zero count. mOTUs were used for modelling if they had a sparsity ≤90% in either birth mode group, indicated by the red lines. This conserves biologically relevant mOTUs that only appear in one group and resulted in 91 of the 959 OTUs being selected.*

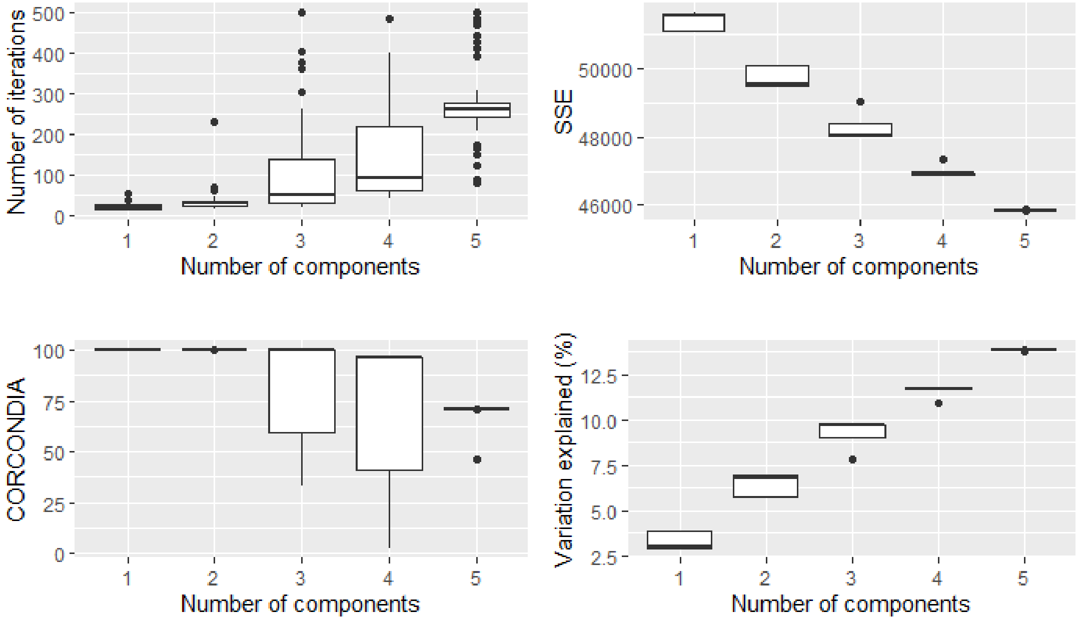


*Supplementary Figure 6: overview plots showing the number of iterations needed, sum of squared error, CORCONDIA score and variance explained for 50 randomly initialized models of 1-5 components each of the processed Shao et al., 2019 dataset. The CORCONDIA scores dropped below 80 at 3-5 components and some of the 3-5 component models did not converge within 500 iterations.*

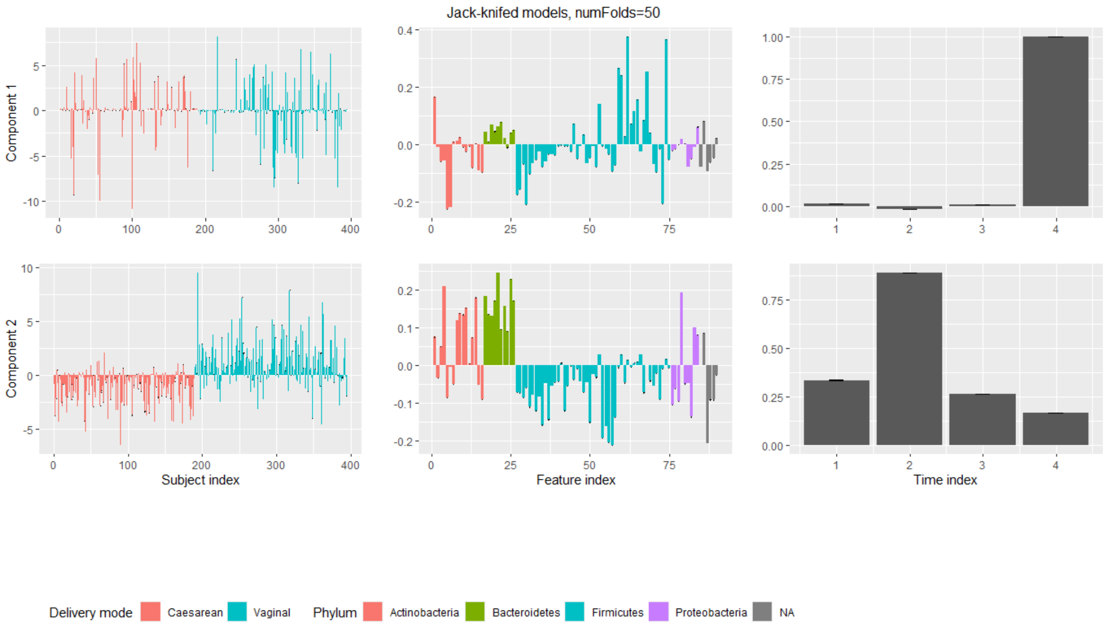


*Supplementary Figure 7 summary of the 50 jack-knifed models of two components created as part of the jack-knifing procedure for the processed Shao et al., 2019 dataset. Modes are shown in the columns (from left to right: subject mode, feature mode, time mode) and components are shown in the rows. Error bars display the range of all values seen for a particular loading across all initialized models. These models show stability in all modes.*

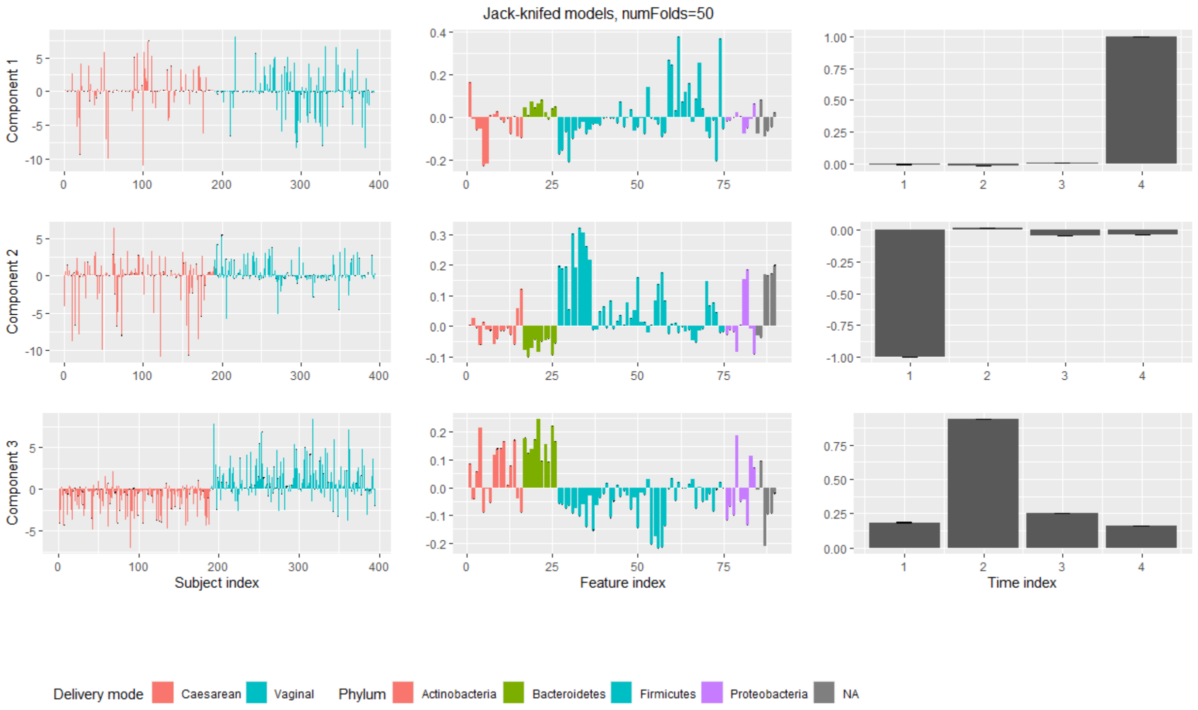


*Supplementary Figure 8: summary of the 50 jack-knifed models of three components created as part of the jack-knifing procedure for the processed Shao et al., 2019 dataset. Modes are shown in the columns (from left to right: subject mode, feature mode, time mode) and components are shown in the rows. Error bars display the range of all values seen for a particular loading across all initialized models. These models show stability in all modes.*

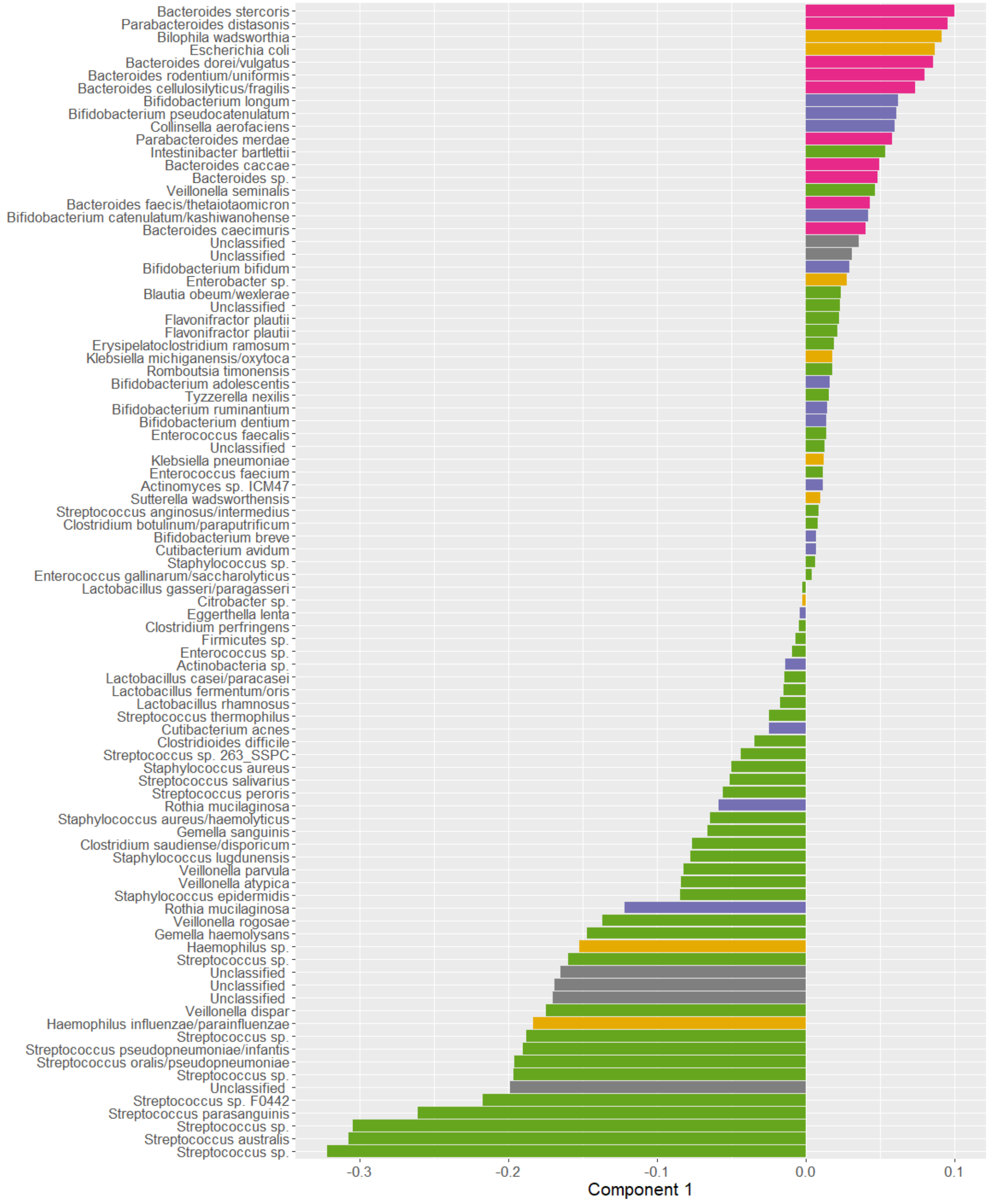


*Supplementary Figure 9: Unfiltered loadings of the microbiota in component 1 of the PARAFAC model of the Shao et al., 2019 dataset. Positively loaded microbiota are associated with vaginally born infants and/or mothers having received antibiotics during labour. Negatively loaded microbiota are associated with caesarean-section born infants and/or mothers not having received antibiotics during labour.*

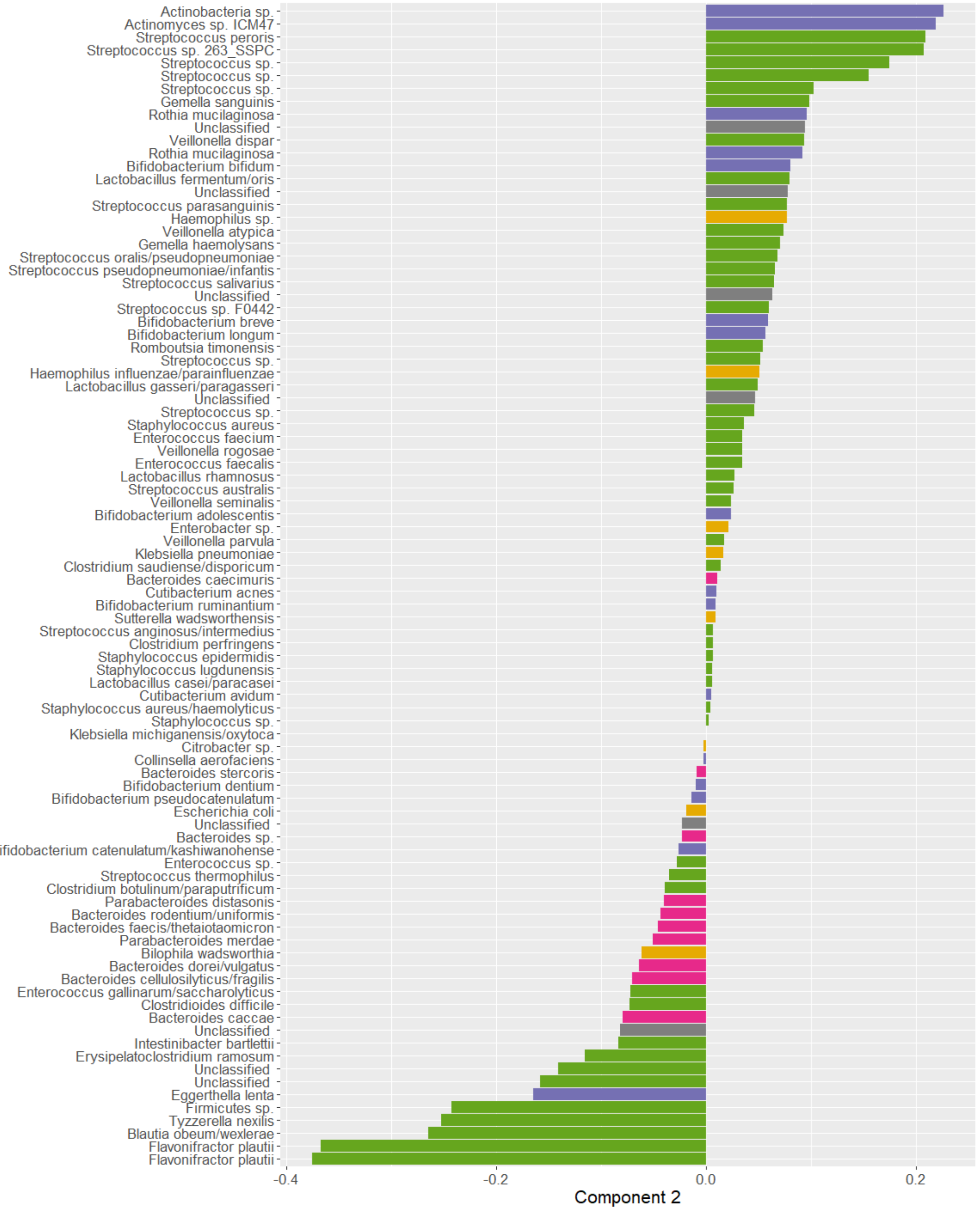


*Supplementary Figure 10: Unfiltered loadings of the microbiota in component 2 of the PARAFAC model of the Shao et al., 2019 dataset. Positively loaded microbiota are associated with vaginally born and/or breast-fed infants. Negatively loaded microbiota are associated with caesarean-section born and/or not breast-fed infants.*

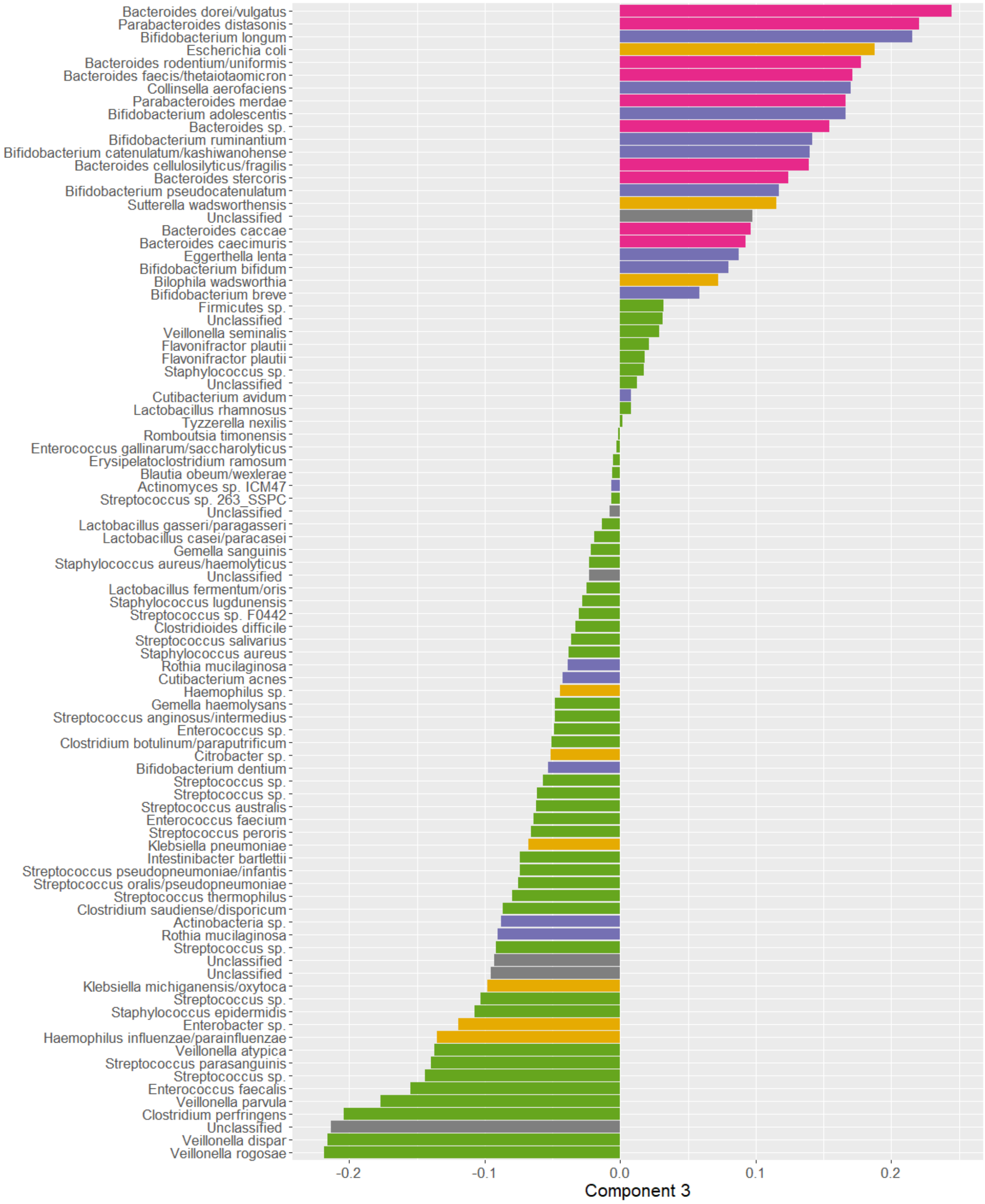


*Supplementary Figure 11: Unfiltered loadings of the microbiota in component 3 of the PARAFAC model of the Shao et al., 2019 dataset. Positively loaded microbiota are associated with vaginally born infants. Negatively loaded microbiota are associated with caesarean-section born infants.*

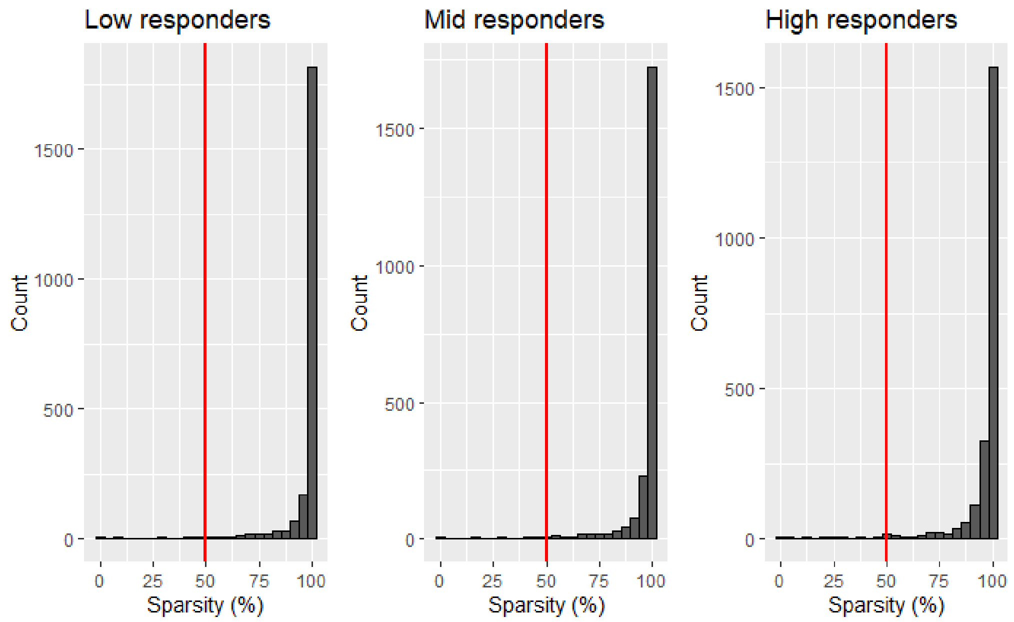


*Supplementary Figure 12: histogram displaying the sparsity of the unprocessed microbial abundance count data for the upper jaw lingual samples in each response group of the van der Ploeg et al., 2024 dataset. Sparsity is defined as the percentage of measurements with a zero count. ASVs were used for modelling if they had a sparsity ≤50% in any response group, indicated by the red lines. This conserves biologically relevant ASVs that only appear in one group and 65 of the 2.253 ASVs remained after this step.*

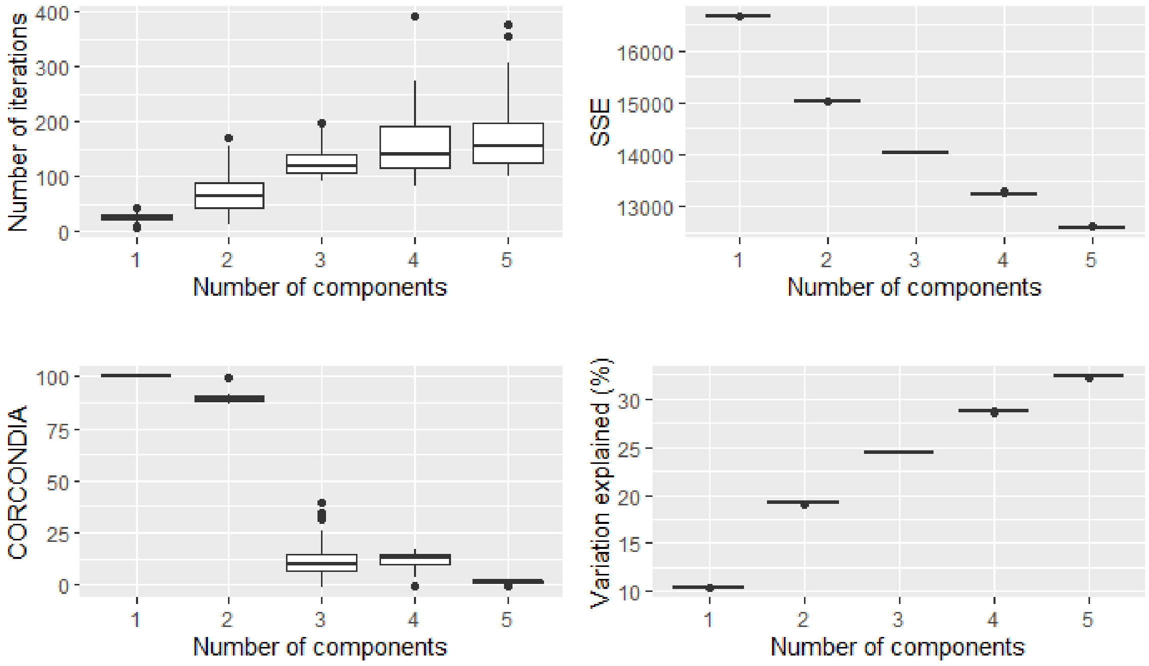


*Supplementary Figure 13: overview plots showing the number of iterations needed, sum of squared error, CORCONDIA score and variance explained for 50 randomly initialized models of 1-5 components each of the processed van der Ploeg et al., 2024 dataset. The CORCONDIA scores dropped below 80 for 3-5 components and the sum of squared errors began to flatten between 2-3 components.*

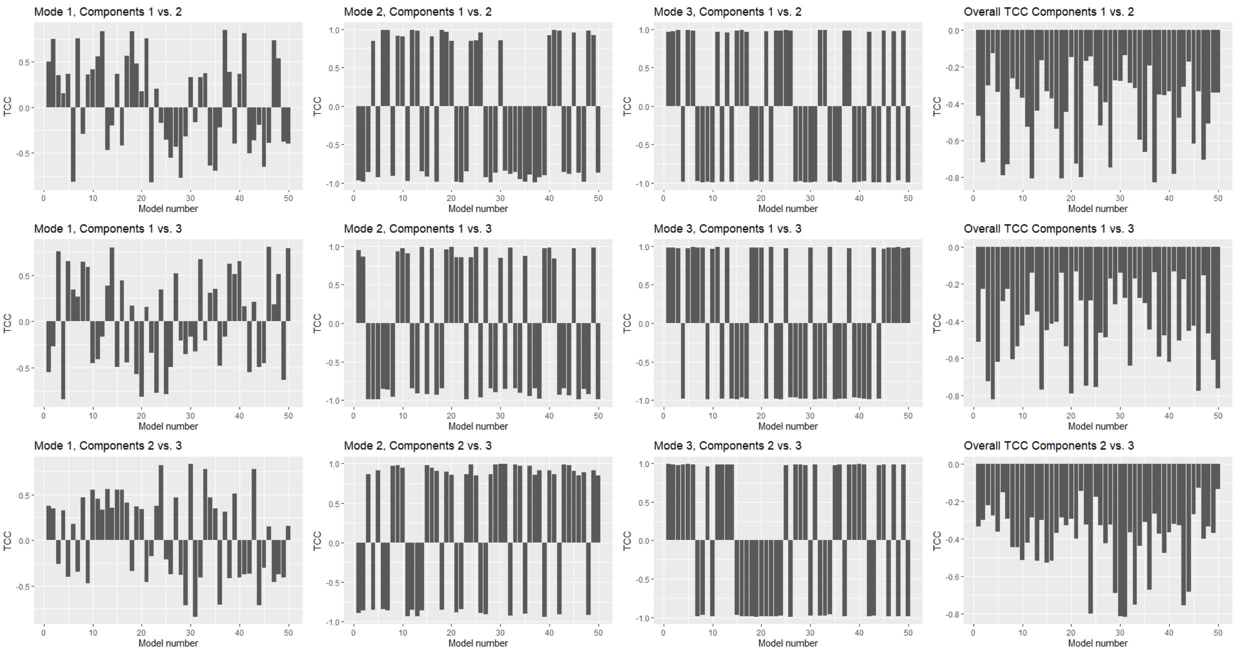


*Supplementary Figure 14: Tucker Congruence Coefficient plot for all 50 randomly initialized models with three components each. The loadings within each mode are compared pairwise in the columns (left: subject mode, middle: feature mode, right: time mode). Comparisons where the absolute value of the Tucker Congruence Coefficient is ≥0.85 leads to the conclusion that the two loading vectors are essentially equal. This is the case for all models with respect to the feature and time mode. Hence the three-component models are likely not suitable for describing the data.*

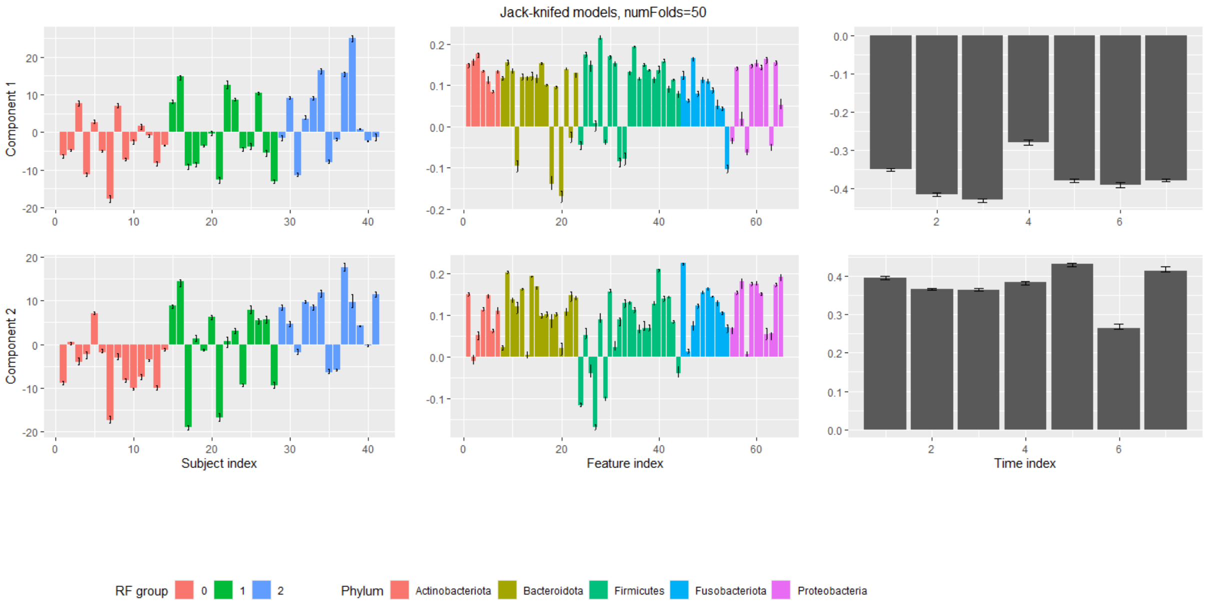


*Supplementary Figure 15: summary of the 50 jack-knifed models of two components created as part of the jack-knifing procedure for the processed van der Ploeg et al., 2024 dataset. Modes are shown in the columns (from left to right: subject mode, feature mode, time mode) and components are shown in the rows. Error bars display the range of all values seen for a particular loading across all initialized models. These models show stability in all modes.*

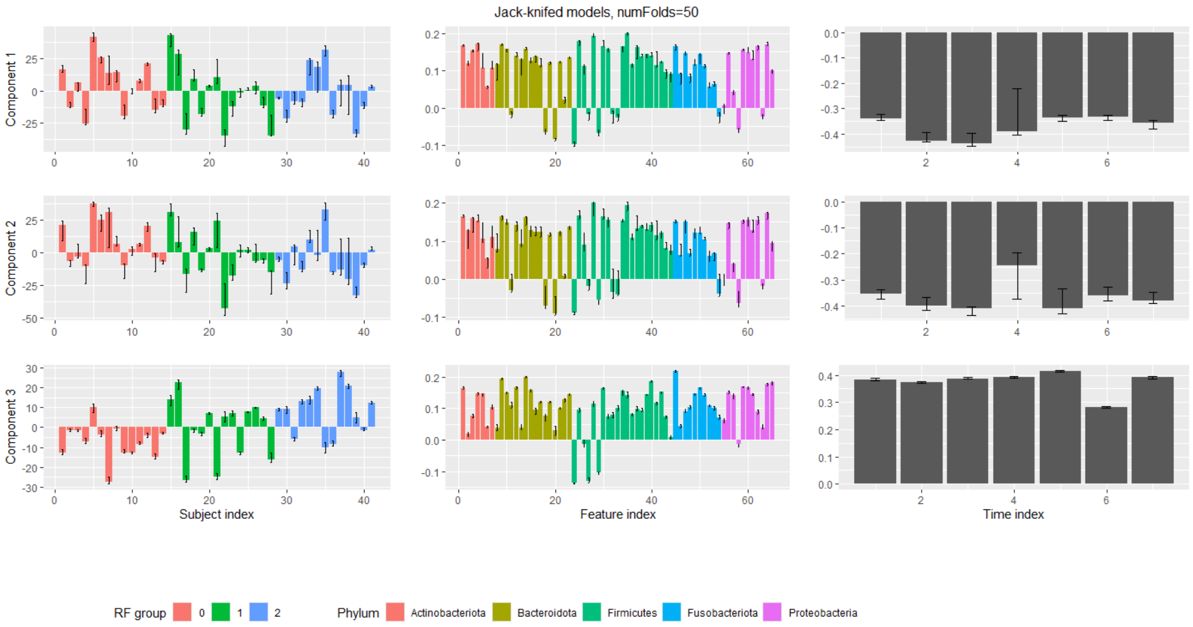


*Supplementary Figure 16: summary of the 50 jack-knifed models of three components created as part of the jack-knifing procedure for the processed van der Ploeg et al., 2024 dataset. Modes are shown in the columns (from left to right: subject mode, feature mode, time mode) and components are shown in the rows. Error bars display the range of all values seen for a particular loading across all initialized models. These models show some instability in the feature and time modes of components 1-2. The third component is largely stable.*

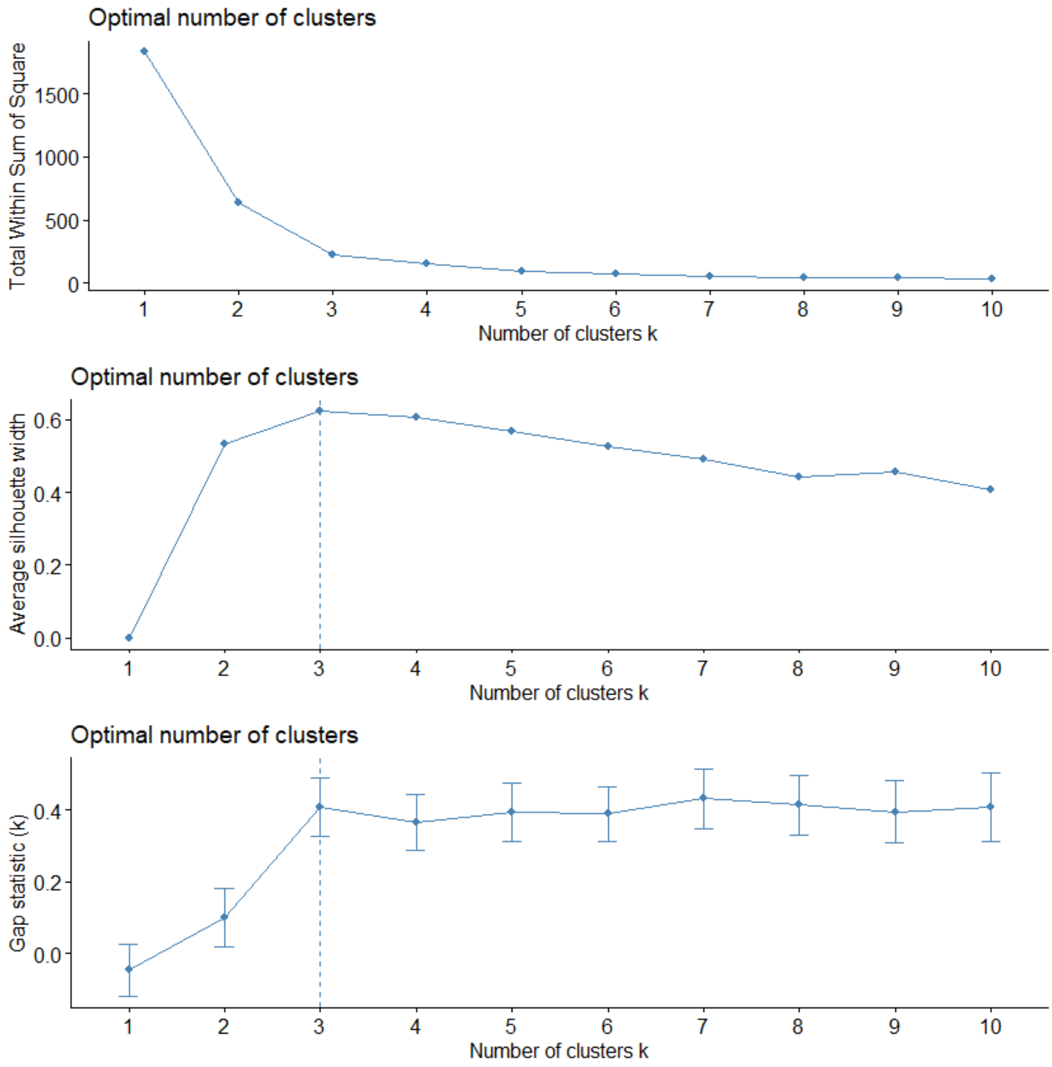


*Supplementary Figure 17: ASV clustering diagnostics for the modelled multi-way array of the van der Ploeg et al., 2024 dataset. The optimal number of clusters is indicated with a dotted line.*

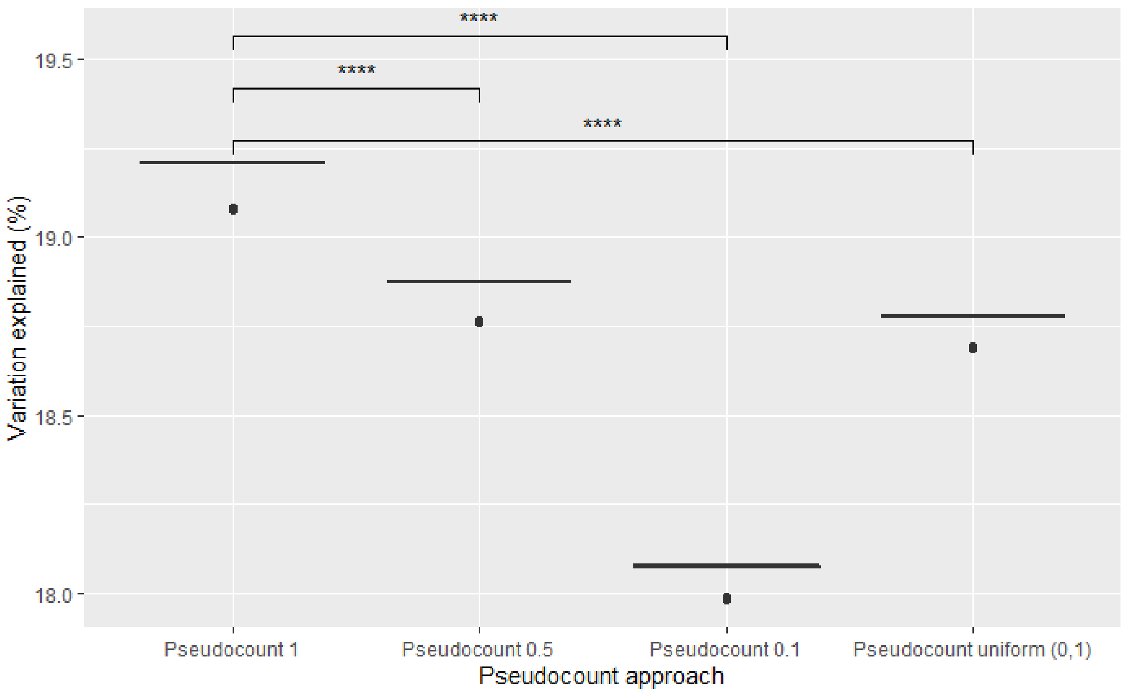


*Supplementary Figure 18: overview of various pseudocount approaches on the eventual PARAFAC model using the van der Ploeg 2024 dataset. Only nonzero elements were used in the calculation of the variation explained. For each method, 50 randomly initialized PARAFAC models were created. Stars indicate p-values corresponding to a Wilcoxon rank sum test between two distributions. (*: p ≤ 0.05, **: p ≤ 0.01, ***: p ≤ 0.001, ****: p ≤ 0.0001)*

### Supplementary Tables

*Supplementary Table 1: The measurements taken per subject in the Shao et al., 2019 dataset. The total number of unique subjects is 395.*

| Time point | Number of subjects with this measurement taken | Number of missing subjects |
|---|---|---|
| 4 | 127 | 268 |
| 7 | 207 | 188 |
| 21 | 136 | 259 |
| Infancy | 122 | 273 |

*Supplementary Table 2: The number of measurements taken per subject in the Shao et al., 2019 dataset. Most individuals have one or more missing measurements. Only four subjects were measured at every time point.*

| Number of measurements | Number of subjects |
|---|---|
| 1 | 237 |
| 2 | 123 |
| 3 | 31 |
| 4 | 4 |

*Supplementary Table 3: Benjamini-Hochberg corrected p-values of Wilcoxon rank-sum tests of various subject metadata available for the Shao et al., 2019 dataset tested against the loadings of the selected PARAFAC model. (*: p≤0.05; **: p≤0.01; *** p≤0.001)*

| Component | Birth mode | Feeding method | Gender | Abx mother during labour | Abx baby in hospital | Abx baby after hospital | Bacteroides profile |
|---|---|---|---|---|---|---|---|
| 1 | **2.4e-4*** ** | 0.14 | 0.65 | **0.0059** ** | 0.74 | 0.95 | **1.5e-4*** ** |
| 2 | **7.3e-9*** ** | **0.024* ** | 0.16 | 0.68 | 0.44 | 0.88 | **7.3e-9*** ** |
| 3 | **1.2e-46*** ** | 0.68 | 0.44 | 0.23 | 0.16 | 0.44 | **9.2e-59*** ** |

*Supplementary Table 4: Benjamini-Hochberg corrected p-values of various subject metadata available for the van der Ploeg et al., 2024 dataset tested against the subject loadings of the selected PARAFAC model. The tests are all Pearson correlation tests, unless specified otherwise. (*: p≤0.05; **: p≤0.01; *** p≤0.001)*

| Component | Plaque% | Bleeding% | RF% | Gender (t.test) | Age |
|---|---|---|---|---|---|
| 1 | **0.029*** | 0.46 | **1.8e-5***** | 0.70 | 0.79 |
| 2 | **0.029*** | 0.37 | **0.029*** | 0.78 | 0.79 |

*Supplementary Table 5: Overview of Benjamini-Hochberg corrected p-values of the permutation test for the differences between the mean sum of relative abundances between the low and high responders per ASV cluster per time point. This is based on 999 permutations of the response groups per time point/ASV cluster combination. (*: p≤0.05; **: p≤0.01; *** p≤0.001)*

| Cluster number | Day -14 | Day 0 | Day 2 | Day 5 | Day 9 | Day 14 | Day 21 |
|---|---|---|---|---|---|---|---|
| 1 | 0.018* | 0.018* | 0.018* | 0.14 | 0.054 | 0.14 | 0.018* |
| 2 | 0.071 | 0.088 | 0.15 | 0.24 | 0.014* | 0.018* | 0.018* |
